# Genomic indicators of risk and resilience in global leatherback turtle populations

**DOI:** 10.64898/2026.05.15.725529

**Authors:** Ekaterina Osipova, Peter H. Dutton, Blair P. Bentley, Sebastian Alvarez-Costes, Katrina F. Phillips, Jamie Adkins, Andrews Agyekumhene, Phil Allman, Ana Rebeca Barragán Rocha, Didiher Chacon-Chaverri, David J. Duffy, Angela Formia, Amy Frey, Alexander R. Gaos, Richard Hamilton, John B. Horne, Shaya Honarvar, Erin L. LaCasella, Deasy Lontoh, Ronel Nel, Anna Ortega, Fitryanti Pakiding, Andhika Prima Prasetyo, Adriana Laura Sarti Martínez, Rotney Piedra-Chacón, Manjula Tiwari, Kelly R. Stewart, João C. A. Thomé, Elizabeth Velez-Carballo, Summer L. Martin, Alana Alexander, Bryan P. Wallace, Lisa M. Komoroske

**Affiliations:** Department of Environmental Conservation, University of Massachusetts, Amherst, MA, USA; Informatics Group, Harvard University, Cambridge, MA, USA; Marine Mammal and Turtle Division, Southwest Fisheries Science Center, National Marine Fisheries Service, National Oceanic and Atmospheric Administration, La Jolla, CA, United States; Department of Biological Sciences, Smith College, Northampton, MA, USA; Department of Anatomy, Faculty of Biomedical Sciences, University of Otago, New Zealand; Dauphin Island Sea Lab, Dauphin Island, AL, USA; Stokes School of Marine and Environmental Sciences, University of South Alabama, Mobile, AL, USA; Department of Marine and Fisheries Sciences, College of Basic and Applied Sciences, University of Ghana, Legon, Ghana; Department of Biology, JN Roth Marine Biology Station, Goshen College, Long Key, FL, USA; Kutzari, Asociación para el Estudio y Conservación de las Tortugas Marinas, México City, México; Latin American Sea Turtles Association/WIDECAST, Costa Rica; The Whitney Laboratory for Marine Bioscience & Sea Turtle Hospital, and Department of Biology, University of Florida, St. Augustine, FL, United States; African Aquatic Conservation Fund, Chilmark, MA, USA; Dipartimento di Biologia, Università degli Studi di Firenze, Firenze, Italy; Protected Species Division, Pacific Islands Fisheries Science Center, National Marine Fisheries Service, National Oceanic and Atmospheric Administration, Honolulu, HI, USA; Asia Pacific Resource Centre, The Nature Conservancy, South Brisbane, Queensland, Australia; School of Mathematical Sciences, Queensland University of Technology, Brisbane, Queensland, Australia; Pacific Cooperative Studies Unit, University of Hawaiʻi at Mānoa, Honolulu, HI, USA; Research and Community Outreach Division, University of Papua, Manokwari, Indonesia; School of Environmental Sciences, Nelson Mandela University, Summerstrand Campus, South Africa; Cooperative Institute for Marine and Atmospheric Research, Honolulu, HI, USA; Research Center for Applied Zoology, National Research and Innovation Agency, Bogor, Indonesia; Comisión Nacional de Áreas Naturales Protegidas (CONANP), 14210 México City, Mexico; Sistema Nacional de Áreas de Conservación, Ministerio del Ambiente y Energía, Heredia, Costa Rica; Ocean Ecology Network, California, USA; St Croix Sea Turtle Project, The Ocean Foundation, Washington, DC 20036, USA; Centro TAMAR-ICMBio, Vitória, ES 29050-335, Brazil; Asociación KUEMAR, Heredia,Costa Rica; Ecolibrium, Inc., Boulder, CO 80303, USA; Department of Ecology and Evolutionary Biology, University of Colorado Boulder, Boulder, CO, 80302 USA

**Keywords:** conservation genomics, genetic load, inbreeding, demographic history, endangered species, sea turtle

## Abstract

Understanding the drivers of genomic health and their consequences for population viability is often overlooked but potentially important to effective conservation amidst the biodiversity crisis of the Anthropocene. Leatherback turtle (*Dermochelys coriacea*) populations have declined globally due to anthropogenic factors, with some populations losing over 90% of their abundance over the past 30-50 years. While conservation efforts have been successful in stabilizing some populations, others continue to decline, and the reasons for these differential trajectories remain unclear. To assess how recent demographic factors, such as population size and decline, influence population genomic health, we combined population monitoring information with medium depth whole-genome and reduced representation resequencing data from globally representative populations. We found that small-stable populations have lower genomic diversity and higher inbreeding than large declining populations, reflecting prolonged small population sizes and limited gene flow. Yet, small-stable populations also show evidence of deleterious allele purging, suggesting genetic resilience. This, combined with lack of detectable genomic erosion over the study period, provides hope for potential recovery of healthy leatherback populations provided that anthropogenic threats are effectively mitigated. However, potential time lags and possible recent increases in inbreeding among close relatives in recently declined populations warrant continued monitoring and assessment. Genomic and abundance-based metrics were less aligned following rapid population declines, emphasizing the different timescales of the evolutionary and demographic processes they reflect, respectively, and the strength in their complementary, integrative use for extinction risk assessments. This also supports that it is not too late to turn the tide for recently declined leatherback populations and that continued investment in conservation efforts and threat reductions are warranted. Collectively, our results highlight how recent and historical demography shapes current genomic health and recovery potential in leatherback turtles, aids understanding of current risks and informs future conservation and management strategies.

## Introduction

As species across the tree of life undergo widespread declines in the Anthropocene ^1^, there is an urgent need to understand the underlying drivers of population declines and obstacles to recovery, which is a key fundamental challenge for conservation. Maintaining genetic diversity within populations to limit inbreeding depression and facilitate adaptive potential has long been a central tenet for conservation ^2,3^, but relationships between genomic health and extinction risk are complex and context-dependent ^4,5^. Theoretical and empirical data from captive populations have provided insight into connections between genomic indicators and fitness impacts ^5,6^, but few studies have validated predictions in wild populations (but see ^7–9^). Thus, it often remains unclear why some threatened species are more sensitive to genetic impacts (e.g., inbreeding depression and/or loss of genetic variation) while others appear resilient ^10^, and the role of genetic factors as possible drivers of decline or impediments to recovery is still rarely considered. Amidst continued calls for effective integration of genomic indicators into extinction risk assessments to complement abundance-based metrics (e.g., population size and trends) ^11–14^, robust understanding of the relationships between demographic and genomic processes is critical.

A central knowledge gap is how the evolutionary and demographic history of a population shapes genetic load, inbreeding, and population resilience. Population declines can result in reduced fitness due to an accumulation of deleterious alleles (genetic load), but long-term small populations can purge strongly deleterious mutations, potentially reducing inbreeding depression (reviewed in ^12^). However, much of how rates of decline, population size, genetic load burdens and purging interact remains an open question for conservation. Empirical studies across taxa show mixed outcomes: in some cases, purging appears effective ^15–17^, whereas in others, substantial genetic load and fitness costs persist despite purging of highly deleterious variants ^7,18^. Moreover, among non-mammalian vertebrates, where almost 20% of all species are currently threatened with extinction (IUCN 2025 ^19^), relationships between historical or recent demography and genetic load are poorly studied, potentially limiting inference to other clades with divergent life history traits and genomic composition.

Leatherback turtles (*Dermochelys coriacea*) provide a compelling system to address this knowledge gap. Their unique physiological, morphological, and ecological adaptations, including long-distance migration, enable them to exploit more wide-ranging marine and thermally complex habitats than any other sea turtle species and most vertebrates ^20,21^. At the same time, several leatherback populations have experienced severe population declines due to anthropogenic factors, such as overharvest of turtles and eggs, accidental mortality via fisheries bycatch, and habitat loss ^22^, as well as changing oceanographic conditions ^23^ that may be exacerbated by climate change ^24^. For example, leatherback abundance in the Pacific Ocean from the mid 1980’s to the early 2000’s decreased by more than 90% ^25–27^. In spite of decades of conservation efforts to protect leatherbacks on nesting beaches and at-sea, depleted leatherback populations have yet to recover in most cases, although a few have shown transient increases or appear to have stabilized at lower abundances ^22,28^. In contrast, similar conservation efforts have generated encouraging success stories for other sea turtle species^29,30^, but the reasons for the stark interspecies differences in responses are unclear. Environmental conditions and resource access can strongly influence population trajectories, and there is some evidence that leatherback turtles may exhibit greater sensitivity to fluctuating environmental conditions ^24,31,32^. In conjunction with differential life history traits relative to other sea turtle species (e.g., reliance on gelatinous prey and pelagic habitats), this could make them more vulnerable to high anthropogenic sources of mortality. However, the extent to which genetic factors like inbreeding depression or low diversity may also influence the extinction risk and recovery potential in leatherback turtles has not been investigated.

Prior work indicates that the low genomic diversity in leatherback turtles is most likely related to long-term, lower population sizes, which could have facilitated purging of deleterious mutations that reduce risks of inbreeding depression ^18,33^, rather than recent loss of diversity from anthropogenic-driven declines ^34^. The very few studies to date investigating relationships among demography, genetic load and inbreeding in non-avian reptiles have found evidence of effective purging in some species but not in others ^35–37^. To our knowledge, there has been no research on purging or other aspects of genomic health (as defined by ^4^) in any turtle species, which are known to have slower evolutionary rates, and distinct life histories relative to other clades ^38,39^. Gaining a robust understanding of how genetic load and inbreeding mechanisms operate and connect to historical and recent demographic factors is therefore key to assessing long-term viability for leatherback turtles specifically, and for informing biodiversity conservation more broadly.

Although leatherback turtles traverse thousands of kilometers across the open ocean, they exhibit strong natal homing ^40^. This innate behavior drives patterns of genetic divergence at much smaller spatial scales relative to their movements ^41^(Fig. 1A). Consequently, there are distinct populations of leatherbacks around the globe with documented differences in demographic trajectories following conservation interventions ^22^, offering a prime opportunity to quantify genomic health metrics and examine their relationships with historical and recent demography and extinction risk. Here, we combine whole-genome and reduced representation resequencing with long-term monitoring data from leatherback populations worldwide to quantify the effects of demographic history on genetic diversity, inbreeding, and genetic load in this long-lived reptile of high conservation concern. We hypothesize that smaller leatherback populations exhibit greater inbreeding and higher genetic load, but that prolonged smaller population size may have enabled more efficient purging of deleterious alleles, potentially reducing realized genetic load. We also predict that genomic indicators best align with demographic classifications when population size and trends have been stable over numerous generations. In contrast, we expect mismatches when there have been rapid changes, reflecting the different timescales of the processes driving abundance-based versus genomic metrics in species with longer generation times. Our findings shed important light on the interplay between demography and genomic health, laying the foundation for incorporating these factors into extinction risk assessments and global sea turtle conservation strategies.

**Figure 1:**
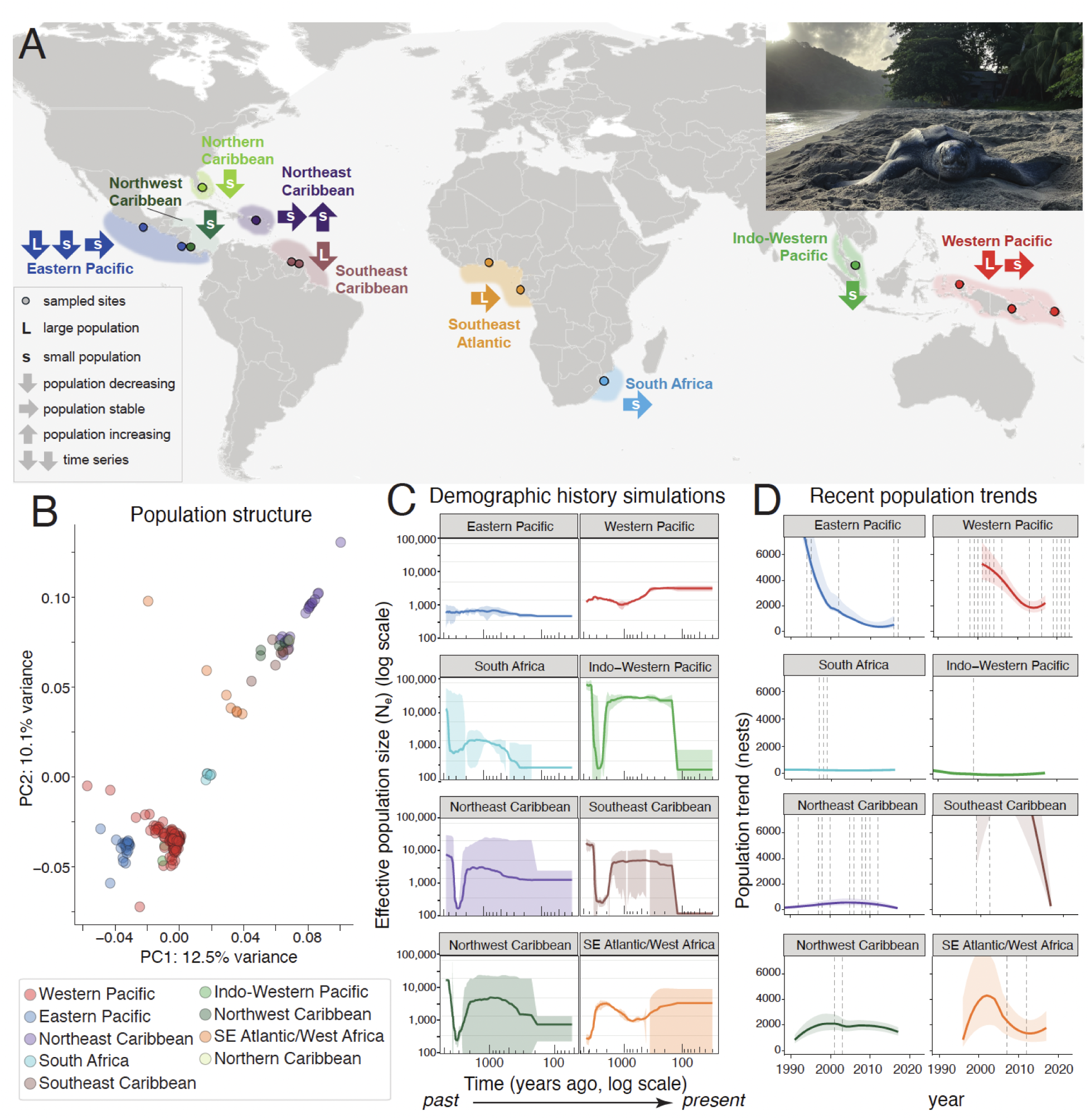
Demographic history and population structure. (A) Geographic distribution of sampled leatherback turtle populations (colored shading) represented by one or more sampled rookery locations (colored points). Arrows depict demographic classifications of the corresponding population for population size (larger or smaller) and trend (increasing, stable or decreasing) at the time of sampling (see Methods for details). Populations with more than one arrow indicate time series data, depicting the population size and trend at each sampling period through time, from left (more historical) to right (more recent). Gray background shading depicts the estimated global geographic range for leatherback turtles. (B) Principal Component Analysis based on whole-genome variation of 222 leatherback sea turtle individuals. (C) Reconstruction of the recent demographic history of leatherback populations using GONE2. The shading represents variance from leave-one-out runs, and the line the full dataset results for each location. (D) Recent population trends estimated from published data collected between years 1967 and 2017 ^48^. All trends were estimated from nest count data, except SE Atlantic/West Africa which used track counts. Vertical dashed lines represent years when samples were collected for genomic analyses. We note that standardized axes facilitate visual comparison, but truncate data for some populations; see Fig. S2B for full unconstrained estimates for all populations.

## Results

### 1. Global population structure and demographic history of leatherback turtle populations

We processed leatherback turtle tissue samples collected from nesting rookeries spanning the entire global distribution range ^42^ (Fig. 1A) to generate whole-genome resequencing (WGR) data for 222 individuals and complementary reduced representation (RAD-capture a.k.a. Rapture) data for 636 individuals from ^43^ (Tables S1,2). These locations are representative of most of the global leatherback populations currently recognized as separate entities for conservation purposes ^42^ (sea turtles are managed under multiple frameworks; see Methods for details). We first confirmed support for using these population designations in downstream analyses by performing population structure analyses on genome-wide single nucleotide polymorphisms (SNPs) from the WGR dataset and a complementary Rapture dataset. Admixture, PCA, and pairwise F_st_ (Table S3) analyses all largely concur with established regional groupings, with clear genetic differentiation among Western Pacific, Eastern Pacific, South Africa, Northwest Atlantic, and Southeast Atlantic populations in one or both datasets (Figs. 1B, S1). Within the Northwest Atlantic locations, both Rapture and WGR datasets detected evidence of finescale genetic structure between the Northeast and Southeast Caribbean (Fig. S1D-E, Table S3,4). Given the distinct demographic trends between these locations ^44^ and that slow mutational rates and long generation times are known to dampen genetic signals of recent isolation in sea turtles ^38^, these results support their retention as separate populations for our analyses. Separation of the Northern and Northwest Caribbean rookeries from other Northwest Atlantic locations was inconclusive due to data limitations (these locations were not represented in the Rapture dataset and had small sample sizes in the WGR dataset; Fig. S1E, Table S4); however, given complementary tagging and other data supporting these locations as distinct demographic units ^44,42^, we also retained them as separate populations for downstream analyses. Separation of the Indo-Western Pacific from the Western Pacific was also supported based on the fine-scale genetic differentiation detected in pairwise F_st_ (Table S3) and ordination analyses (Fig. 1C, S1D-E, Table S4), consistent with separate results from microsatellite nDNA analysis ^45^. Taken together, our population structure results generally correspond with existing population management units with minor differences (see Methods for details) to group nesting rookeries effectively for analyzing demographically independent populations.

We then evaluated ancient and contemporary leatherback effective population sizes (Ne)^46,47^ with two complementary methods. Using PSMC, all populations exhibit similar ancestral demographic histories and historical (over 20,000 years ago) lower effective population sizes (Fig. S2A), consistent with prior studies ^34^. Since contemporary Ne is generally of more relevance to conservation compared to historical demographic estimates, we also examined linkage disequilibrium (LD)-based Ne estimates that have greater ability to accurately estimate more recent demography and examine differences between populations over the last 4,000 years (∼150 generations). All populations except the Eastern Pacific and Western Pacific exhibit evidence of a bottleneck and recovery around 2,000 years ago. Over the past 1,000 years, some populations displayed relatively consistent Ne estimates, such as the consistent smaller Ne in Eastern Pacific and South Africa populations (Fig. 1C). In contrast, other populations exhibited pronounced fluctuations, including the Southeast Caribbean and Indo-Western Pacific populations that showed steep and abrupt Ne declines within approximately the last 100 years.

### 2. Lower heterozygosity and higher inbreeding in small populations

Following population structure characterization, we used published abundance-based demographic data ^22,42,44,49,50^ to classify each WGR sample by population size (large or small) and trend (declining, stable, or recovering) at the time of sampling for each population (Fig. 1A,D). Notably, with the exception of the Southeast Atlantic population, leatherback populations are now smaller than they were at the time of sampling (Fig. 1D). For example, the Western and Eastern Pacific populations were initially sampled when they were larger prior to or during recent declines, whereas others have remained consistently smaller over time (e.g., South Africa and Northeast Caribbean populations). We then defined five groups based on combinations of size and trend: larger-stable, larger-declining, small-stable, small-declining, and small-recovering (Fig. 1A; see Methods for details). We conducted quantitative statistical comparisons between only the larger-declining and small-stable groups because they had sufficient sample sizes, and qualitative assessments of observed patterns among other groups with insufficient sample sizes.

We found that larger-declining populations have significantly higher genome-wide heterozygosity compared to the small-stable ones (p = 1.0e-6; Fig. 2B), despite similar total SNP density (p > 0.05; Fig. 2A,B). Regardless of demographic trend, small populations generally exhibit lower heterozygosity (Fig. S3B). However, samples from the Indo-Western Pacific and the Western Pacific populations that were collected when these populations became small following rapid declines (Fig. 1D, Fig. S2B) were exceptions to this pattern, displaying higher heterozygosity similar to that of larger-declining populations (Fig. S3). Conversely, the larger-stable population in the Southeast Atlantic has unexpectedly low heterozygosity, comparable to that of small populations. These results demonstrate the significant effects of population history and genetic background on genomic diversity, likely reflecting processes that occur over a longer timescale relative to when samples were taken.

**Figure 2:**
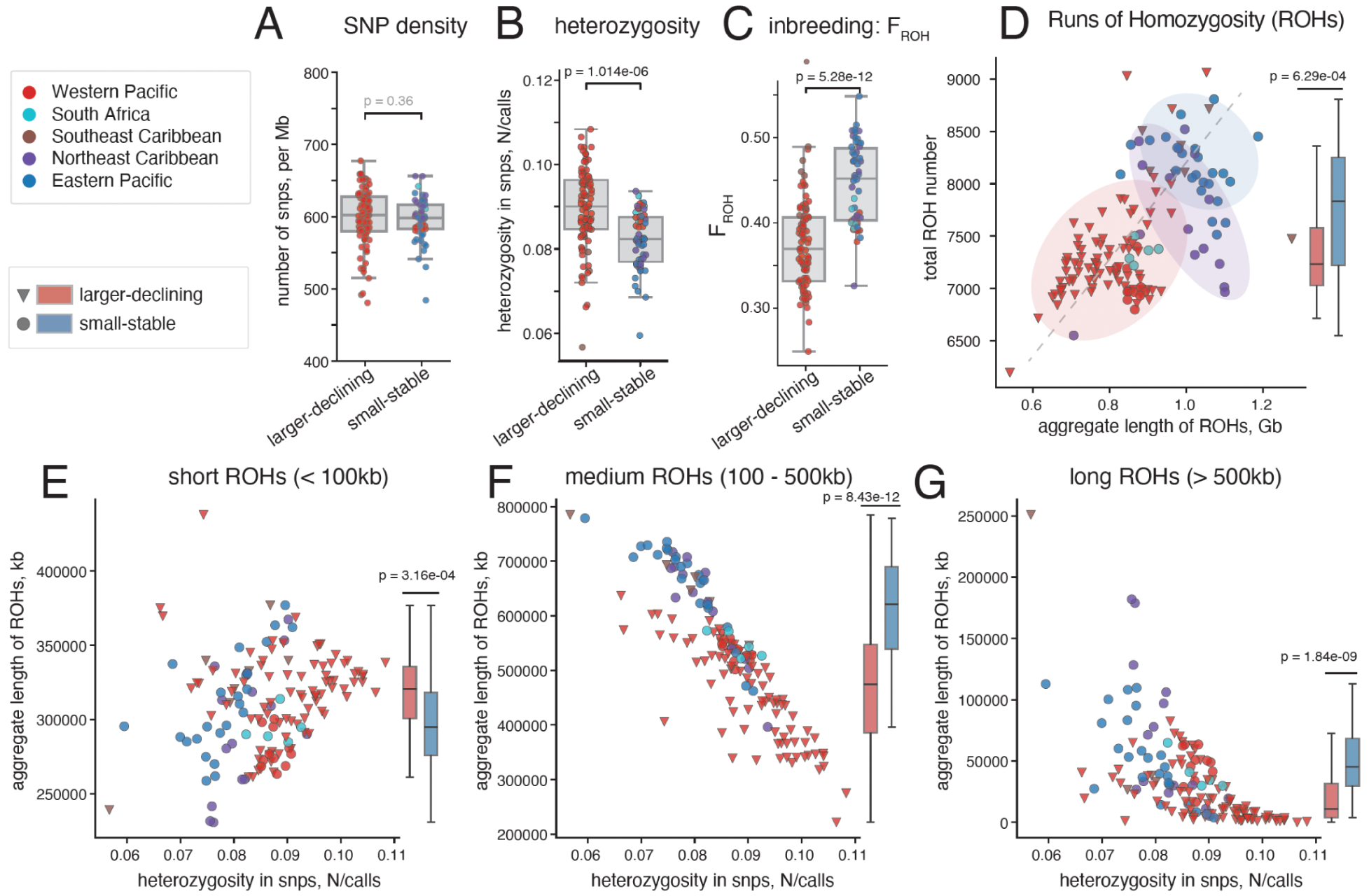
Heterozygosity and inbreeding. (A) There is no difference in single nucleotide polymorphism (SNP) density between larger-declining and small-stable populations. (B) Small-stable populations have lower genome-wide heterozygosity (calculated per called SNP) than larger-declining populations. (C) Inbreeding is higher in small-stable populations compared to larger-declining populations (F_ROH_ designates the inbreeding coefficient). (D) Total number of runs of homozygosity (ROHs) and their aggregate length. Shaded areas highlight where the majority of corresponding samples from the Western Pacific, Northeast Caribbean, and Eastern Pacific populations lie. (E-G) Total length of the genome contained in ROHs of different sizes: shorter than 100kb (E), between 100 and 500kb (F), and longer than 500kb (G). For statistical comparisons between these two groups, a two-sided Mann-Whitney U test was used (all panels). Boxplots shown on panels D-G refer to the y-variable on the corresponding plot. P-values > 0.05 are presented in grey.

To assess inbreeding in leatherback populations, for each individual, we quantified runs of homozygosity (ROHs), i.e., long stretches of DNA where an individual has inherited identical genetic segments from both parents ^18,37^. Consistent with their lower heterozygosity, small-stable populations have a higher fraction of their genome within ROHs than larger-declining ones (F_ROH_, p = 5.3e-12; Fig. 2C). Small-stable populations also show higher numbers of ROHs than larger-declining populations (Fig. 2D), consistent with expectations based on effective population size ^51^. Populations in other demographic groups displayed patterns concordant with heterozygosity results, including the large-stable Southeast Atlantic population generally showing ROH patterns similar to small-stable populations, and the small-declining Indo-Western Pacific population resembling larger-declining populations (Fig. S4); however, robust inference is limited by the sample sizes of these populations.

To better understand the timing of inbreeding and other recent population history events, we classified ROHs by length into short (<100 kb), medium (100-500 kb), and long (>500 kb) categories, based on parameter testing and leatherback turtle recombination rates (see Methods). The number of medium-length ROHs contributes the most to the variation between populations (Figs. S4A, S5) and is correlated with heterozygosity (Figs. 2F, S6), reinforcing the conclusion that population history influences contemporary genetic diversity levels. While larger-declining populations have slightly more short ROHs (p = 3.2e-4; Fig. 2E-G), small-stable populations have significantly more medium and long ROHs (p = 8.4e-12 and p = 1.8e-9, respectively; Fig. 2E-G). The presence of these long ROHs suggests insufficient time for recombination to break these regions into shorter segments, further supporting a greater prevalence of recent inbreeding in small populations.

Notably, some individuals in the Northeast Caribbean population have a relatively small total number of ROHs, similar to those observed in larger populations (Fig. 2D). However, the aggregate length of the ROHs in these individuals is very high, comparable to that of small populations (Fig. 2D), which is explained by a predominance of long ROHs (>500kb; Figs. 2G, S4A). This pattern suggests recent inbreeding between close relatives ^51^. We did not observe a correlation between this signal and the time when the samples were collected (Fig. S6A), suggesting that the inbreeding between closely related individuals is variable.

### 3. Evidence of purging of deleterious alleles in small populations

Genetic drift in small populations may overwhelm purifying selection, allowing high-impact alleles, such as frameshifting mutations and in-frame stop codons, to increase in frequency or become fixed over time ^12,52^. In some cases, elevated inbreeding may facilitate the purging of these deleterious - or high-impact - alleles, thereby providing some resilience against the impacts of inbreeding depression and improving long-term population viability ^16,18,37,53^. In other cases, inbreeding increases the expression of harmful recessive mutations without effective purging, leading to a higher mutation load and an increased risk of mutational meltdown ^37,53,54^. Here, we tested whether higher levels of inbreeding in small populations leads to the accumulation of mutation load or if putatively harmful alleles are being effectively purged.

To assess mutation load in leatherback populations, we annotated variants based on their putative fitness effects using snpEff, classifying them into high-, moderate-, and low-impact categories. High-impact variants typically disrupt gene function by altering the reading frame and are therefore more likely to have strong deleterious consequences for fitness. As strongly deleterious variants have greater effects on fitness, evidence of purging should be most pronounced in this category ^12,55^. We found that small-stable populations show a similar number of high-impact variants (p = 0.22), but carry a higher fraction of moderate- and low-impact variants than do larger-declining populations (p = 0.0025 and p = 0.0014, respectively; Fig. 3A), suggesting higher total mutation load in small populations, but with potential purging of strongly deleterious mutations (Fig. 3A). Other demographic groups (small-declining, larger-stable and small-recovering) qualitatively exhibited similar fractions of high impact variants to the larger-declining and small-stable groups, however small sample size and substantial variation in moderate and low impact variants among individuals limits these comparisons (Fig. S7).

**Figure 3:**
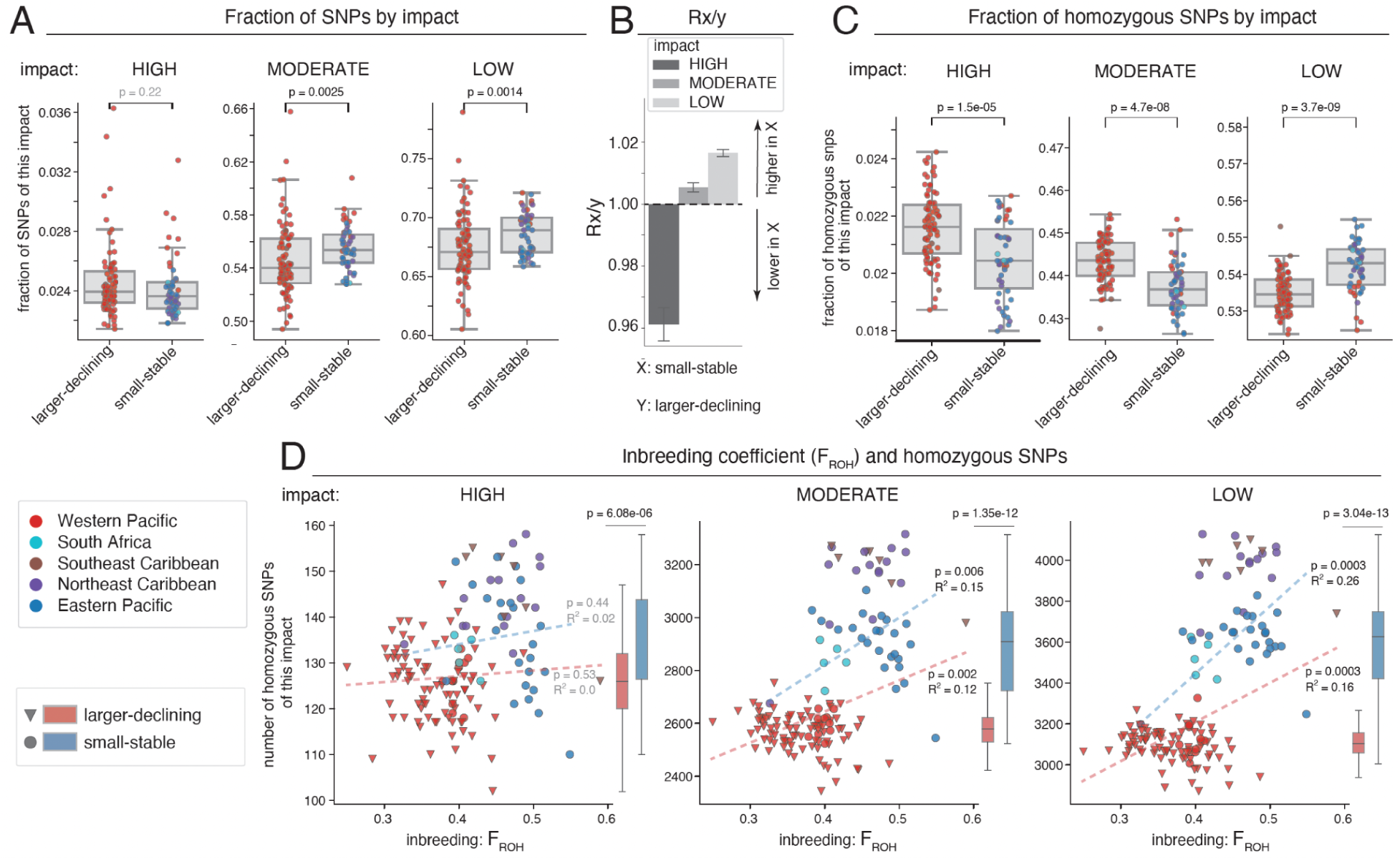
Genetic load and purging. (A) Total number of single nucleotide polymorphisms (SNPs) by putative fitness impact shows a higher fraction of moderate and low impact SNPs in small-stable populations compared to larger-declining populations. (B) Rxy analysis shows a higher mutation load of moderate and low-impact variants in small-stable populations compared to larger-declining populations. Error bars represent standard error from block jackknife approach. (C) The fraction of high and moderate-impact homozygous SNPs is lower in small-stable populations compared to larger-declining populations. (D) The higher inbreeding in small-stable populations (F_ROH_ designates the inbreeding coefficient) is positively correlated with the total number of low- and moderate-impact, but not high-impact homozygous SNPs. R^2^ values and p-values reflect the results of regression analysis for each group. Boxplots shown on panel D refer to the y-variable on the corresponding plot. For statistical comparisons between these two groups, a two-sided Mann-Whitney U test was used. The error bars in boxplots (A,C,D) represent 95% confidence intervals. P-values > 0.05 are presented in grey.

These observations were further supported by Rxy analysis, which compares the relative frequency of alleles of different impacts between populations ^56^. An Rxy value above 1 indicates that a given variant class is more common in the target population (here, small-stable) than in the comparison population (here, larger-declining). For moderate- and low-impact variants, we found that Rxy values are above 1, indicating a small but significant higher mutation load of these deleterious variants in small-stable populations. In contrast, Rxy for high-impact variants are significantly below 1 with a substantially larger effect size (Fig. 3B), confirming that strongly deleterious alleles are less frequent in small-stable populations. Together, these results point to a pattern that, relative to larger-declining populations, small-stable populations have an elevated mutation load overall, coupled with effective purging of the most harmful alleles.

While total variant counts reflect the total mutation load, the fraction of homozygous deleterious alleles measures the realized genetic load more directly. If purging takes place, it is most effective when deleterious alleles become homozygous and are exposed to selection ^12,16,54^. Thus, to further test for evidence of purging, we quantified the fraction of homozygous variants by impact. Small-stable populations exhibit significantly higher fractions of low-impact homozygotes (p = 3.7e-9) but significantly lower fractions of high-impact homozygotes (p = 1.5e-5; Figs. 3C, S8A). Notably, we also detected a lower fraction of moderate-impact variants (p = 4.7e-08; Figs. 3C, S8A), suggesting that small-stable leatherback populations can effectively purge not only the most deleterious mutations but also moderately deleterious ones. These results were independently confirmed using an alternative variant effect annotation method, ANNOVAR, that shows a significantly higher fraction of high-impact homozygotes (p = 1.7e-8; Fig. S8B) and a lower fraction of moderate-impact homozygotes (moderate: p = 0.025; high: not significant with p = 0.097) in small-stable populations compared to larger-declining ones. Collectively, these analyses provide support for the purging of deleterious alleles in small-stable populations relative to larger-declining populations. The other demographic groups qualitatively were most similar to the patterns of the small-stable category; however, again, the limited sample size precludes robust inference for these groups (Figs. S3B, S8).

To further assess the interplay between realized genetic load and inbreeding in the context of possible inbreeding depression, we examined the relationship between the inbreeding coefficient (F_ROH_) and the number of homozygous variants by putative fitness impact. We found that higher inbreeding in small-stable populations is associated with an increased total number of low- and moderate-impact homozygotes (R^2^ = 0.26 with p = 0.0003 and R^2^ = 0.15 with p = 0.006, respectively; Fig. 3D, Table S5). However, inbreeding had no significant correlation with the total number of the most deleterious, high-impact homozygotes (R^2^ = 0.02 with p = 0.44) (Figs. 3D, S9). This pattern further supports the conclusion that while strongly deleterious alleles are being purged in small-stable populations, less harmful mutations are more likely to reach high frequencies or fixation, and be retained in homozygous states due to inbreeding between close relatives. Notably, the Northeast Caribbean population exhibits the highest total number of low- and moderate-impact homozygotes, suggesting that the recent inbreeding has likely contributed to the accumulation of deleterious alleles in this population.

### 4. No evidence of genomic erosion over recent decades

Genetic changes, such as erosion of genomic diversity or shifts in inbreeding levels or genetic load, can require several generations to become evident at the genomic level ^54^. For two populations - the Eastern Pacific (N=39) and Northeast Caribbean (N=28) - we used a time series of leatherback samples collected across several decades from 1995 to 2020. This provided a unique opportunity to examine ongoing changes in the genomic health of these populations. We used information on life stage - adult or hatchling - and year of collection to assign samples to three consecutive generations, and then tested for trends using generalized linear models (see Methods for details). We found no significant changes in heterozygosity, total inbreeding, or highly deleterious realized genetic load in either population between generations (Fig. 4). In the Eastern Pacific population, we detected a potential increase in medium and long Runs of Homozygosity (ROHs) (FDR-corrected p = 0.02 and p = 0.02 respectively, Fig. S10A, Table S6), accompanied by a marginally significant decrease in short ROHs (p = 0.06, Fig. S10A). This pattern suggests some mating between close relatives in this population in recent generations. In the Northeast Caribbean population, we observed significant decreases in short ROHs (p = 0.01) as well as moderate (p = 0.05) and low impact homozygous variants (p = 0.04), but a lack of significant changes in long ROH or high impact variants (Fig. S10A,B). However, the substantial variation in ROH and genetic load metrics indicates that more samples are needed for robust data interpretation. Collectively, our results indicate that over approximately 25 years, there has been little change in the overall genomic health of these two focal populations, but the observed increases in recent inbreeding events warrant continued monitoring to detect and mitigate possible future genomic erosion and inbreeding depression.

**Figure 4:**
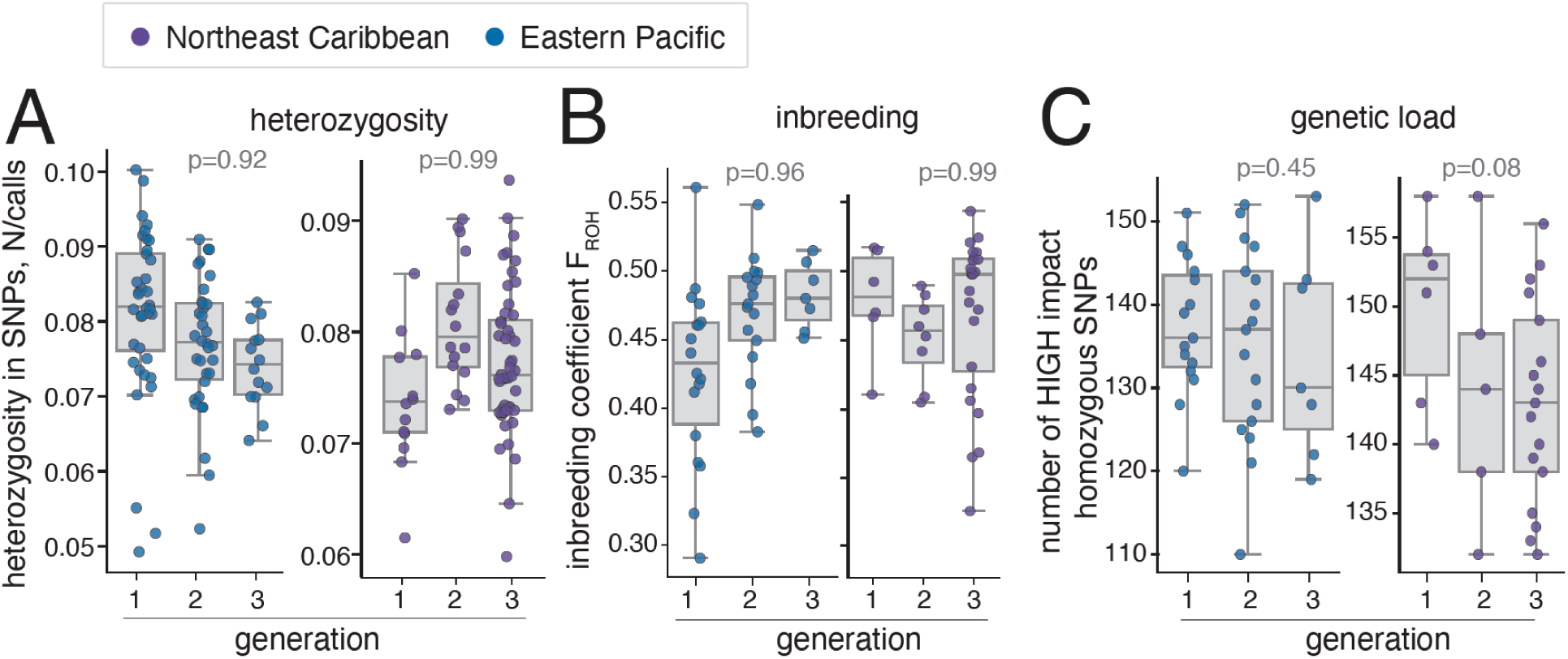
Temporal analysis of genomic health in leatherbacks. (A-C) Temporal analysis of heterozygosity (A), inbreeding measured by inbreeding coefficient F_ROH_ (B), and genetic load measured by the number of high-impact homozygotes (C) in samples collected in the Eastern Pacific and Northeast Caribbean across three generations does not show any significant trends (using threshold for FDR-corrected p < 0.05). To test trends across generations, generalized linear models were used (see Methods for details).

### 5. Structural variation

Inversions and other structural variants are increasingly being associated with local adaptation ^57^ and genomic incompatibilities ^58^. Inverted regions of the genome have the potential to act as ‘super genes’, linking together adaptive alleles that provide a fitness advantage under specific conditions ^59^. Therefore, characterizing structural variation is relevant for a variety of conservation contexts, such as preservation or loss of adaptive variation for key traits in declining populations and avoiding the introduction of maladaptive variation during population supplementation or genetic rescue ^60,61^. To identify structural variants that could potentially contribute to local adaptation or genomic incompatibilities in leatherback sea turtle populations, we ran local windowed principal component analysis (PCA). We identified 52 distinct regions of local population differentiation across the leatherback genome (Table S7), including two putative segregating inversions at the end of chromosome 14 (Fig. 5) containing 35 zinc-finger protein genes, nine immune-related genes, including seven copies of butyrophilin subfamily genes, two L-amino-acid oxidase genes, and two olfactory receptor genes. Additionally, we found copies of genes tied to tumorigenesis (targeting protein for Xklp2 and E3 ubiquitin-protein ligases) within these regions, as well as genes encoding histones, rRNAs, and tRNAs. This region of chromosome 14 has been previously associated with lower copy numbers of olfactory receptor, zinc-finger protein, and immune genes (including the major histocompatibility complex; MHC) in leatherback turtles compared to the hard-shelled species ^34,62^, supporting that its evolution and genomic variation play important roles in immunological and ecological adaptation within and among sea turtle species. We also identified another region of interest on chromosome 6 (Fig. 5E), which shows local structural differentiation between a number of the Eastern Pacific individuals and the rest of the samples. Six of eight genes annotated in this region encode long non-coding RNAs with potential regulatory functions. The other two genes, RAG1 and RAG2, encode proteins that activate immunoglobulin V(D)J recombination, and are part of a previously identified larger *Rag* gene cluster in other jawed vertebrates ^63^. These are essential immune genes that reshuffle DNA segments in developing B and T immune cells to create unique antibodies and T-cells, allowing them to recognize foreign pathogens. The identified putative structural variants could contribute to local immune adaptations in leatherback populations.

**Figure 5.**
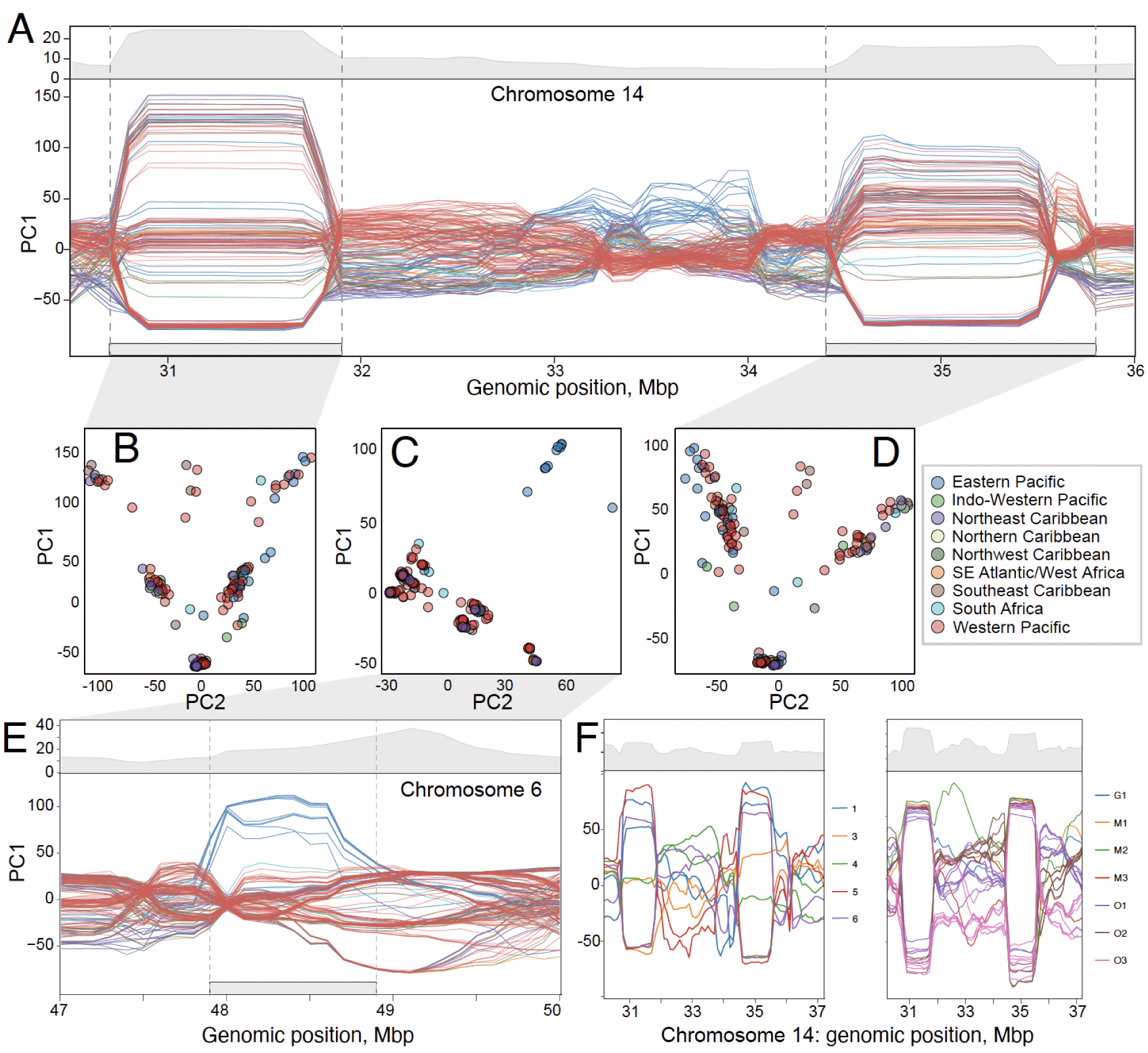
Putative structural variation in leatherback genomes. Sliding window Principal Component Analysis (PCA) for the end of chromosome 14 (A) and chromosome 6 (E); lines are colored by population, and grey shaded plots above the line plots show percent of variation explained by PC1. Putative inversions were identified by splitting of samples into three distinct groups representing the homozygous dominant and recessive, and heterozygous genotypes (see Methods for details). Associated PCA plots of PC1 and PC2 for the first (B) and second (D) putative inversions on chromosome 14, and the region of interest on chromosome 6 (C). Mbp = 1,000,000 base pairs. (F) Putative inversions region on chromosome 14 in samples from known related individuals in the East Pacific population (left, numbers refer to 6 mother-hatchling pairs) and the Northeast Caribbean population (right, G1=grandmother, M1-3 = mother, O1-3=hatchling offspring of mothers with corresponding mothers), depicting homozygous and heterozygous individuals (see text). Note that in the left panel all three genotypes are present (homozygous reference, heterozygous, and homozygous alternate), while in the right panel individuals only have two of the three possible genotypes.

Possible genomic incompatibilities between individuals with larger structural rearrangements is a potential concern for conservation programs, especially in the context of selecting individuals for genetic rescue or captive breeding programs. We leveraged a subset of samples in our dataset of known parent-offspring relationships from the East Pacific and Northeast Caribbean populations to explore this question. In the East Pacific a heterozygous mother produced a homozygous hatchling and vice versa: homozygous mother - heterozygous hatchling (Fig. 5F). In the Northeast Caribbean one grandmother and three mothers were all homozygous in the chromosome 14 inversion region, but all three mothers produced heterozygous hatchlings (Fig. 5F). More work is needed to fully understand the genomic compatibility of these structural variants, but these preliminary results suggest that heterozygous individuals are able to successfully complete embryonic development, reach maturity and reproduce.

## Discussion

Understanding relationships between demographic history and genomic health indicators and how they impact population viability is essential for their effective use in biodiversity conservation. By integrating whole-genome resequencing with demographic data across leatherback populations with divergent sizes and demographic trajectories, we demonstrate that longer-term evolutionary history plays the dominant role in shaping current patterns of genetic diversity, inbreeding, and mutational load. Specifically, small-stable populations show lower genome-wide heterozygosity and higher inbreeding compared to larger-declining populations. Despite this reduced diversity, small-stable populations show clear evidence of effective purging of highly deleterious mutations, aligning with recent studies in other long-lived vertebrates where persistent small effective population size allowed purifying selection to remove the most harmful mutations over evolutionary timescales ^16,18,55^. Coupled with little genomic erosion detected to date in recent temporal samples, this suggests some degree of resilience against inbreeding depression and provides hope for leatherback populations as long as pervasive threats, such as bycatch, harvest, and habitat loss, can be effectively mitigated. We also show that abundance-based and genomic population health indicators are less aligned following rapid demographic changes, emphasizing the influence of population history and other factors on these metrics and the strength in their complementary, integrative use for extinction risk assessments ^4,13^.

We found that moderately deleterious mutations are purged less efficiently than highly deleterious ones. Compared to larger-declining populations, small-stable populations show fewer moderate-impact homozygotes, indicating some purging of these, yet Rxy analysis still suggests accumulation of these variants. This discrepancy implies that purging efficiency depends on the strength of mutation effects on fitness: strongly deleterious alleles are effectively eliminated, whereas low- to moderately deleterious variants can drift to higher frequencies, contributing to long-term mutation load. Despite purging, the overall higher genetic load and accumulation of mildly deleterious variants in small-stable populations could impact fitness and future resilience of these populations, similar to ibexes or cranes ^64,65^. Recent work has also highlighted that purging does not preclude inbreeding depression impacts on population viability ^3,7,18^. Additionally, the combination of high overall genetic load and fewer highly deleterious homozygotes indicates that small-stable populations harbor more harmful alleles in the heterozygous state. Although these alleles may not have immediate fitness consequences, they could become exposed under sustained inbreeding, threatening population viability. Thus, even with evidence of purging and apparent genetic resilience, high genetic load and accumulated mildly deleterious variants could still impair fitness such that inbreeding depression may remain a risk for leatherback turtles, especially if rapid declines of historically larger populations outpace the time needed for effective purging ^5^.

In addition to observing a clear effect of population size on genomic signatures, contemporary LD-based Ne estimates were generally aligned with recent demography, including detecting some signatures of demographic declines in the last several generations (e.g., Indo-Western Pacific and Southeast Caribbean; Fig. 1C, Fig. S2B). However, our results also show that genomic health and abundance-based indicators of extinction risk are often poorly aligned following rapid demographic change. For example, we qualitatively observed that within the Western Pacific and the Indo-Western Pacific populations, individuals from locations with small population sizes following recent, rapid declines, display heterozygosity, inbreeding, and genetic load patterns more similar to the larger-declining populations. In fact, the samples analyzed here from the Indo-Western Pacific population were collected from some of the last nests that emerged in peninsular Malaysia before the rookery was extirpated ^66^. This indicates that when declines are rapid, genomic diversity can remain higher even when extinction is imminent (as found in ^8^), and that shared genetic ancestry as well as population-specific history contribute to patterns of genomic health indicators. This pattern is consistent with theory ^3,12^ and previous work underscoring the need to consider demographic history alongside abundance-based indicators when assessing extinction risk ^4,11^, and is likely particularly important for species with long generation times and slower evolutionary rates like sea turtles. However, we acknowledge that the unequal sample sizes across all populations and categories limit our ability to fully tease apart the effects of population size and trend, and we hope that future studies can fill these gaps.

Given the rapid rates of demographic change, it is not surprising that samples collected before and after recent strong declines (abundance reduced by >90% in some cases) do not yet show bottleneck signatures ^3,10^. Nonetheless, the lack of evidence for genomic erosion in temporal analyses of East Pacific and Northeast Caribbean populations suggests some genomic resilience to recent anthropogenic impacts, and could be encouraging for recovery if other threats can be mitigated. However, long generation times may extend lags between demographic change and detectable genomic signals ^3,12^, so that even decades of monitoring may be insufficient to fully capture emerging genetic risks and/or that more sensitive metrics such as estimating loss of rare alleles may be needed ^67^. Recent studies have suggested that inbreeding metrics are better predictors of inbreeding depression and fitness relative to other metrics of genomic health^68,13^, including evidence that long ROH burden has the strongest effect on inbreeding depression and negative fitness consequences ^13^. Although appropriate ROH thresholds vary across taxonomic groups due to the influence of recombination rates ^13^ and methodological choices can influence ROH detection ^69^, our finding that long ROH burdens in leatherbacks were substantially lower relative to what has been observed in other inbred species suggests that most leatherback populations are still large enough to prevent widespread contemporary inbreeding. Such resilience may be at least partially driven by sea turtle life history traits; in contrast to mammals and birds (the taxa in which majority of inbreeding depression studies have been conducted) that typically have few offspring and parental care, sea turtles can produce many hundreds of hatchlings over numerous egg clutches with high rates of multiple paternity ^70,71^, and be highly iteroparous over their lifetimes. These traits may buffer against mating between close relatives, and potentially facilitate purging of deleterious variants because inbreeding depression can impair embryonic development or fitness of some hatchlings, but with high numbers of genetically different offspring it is more likely that some will inherit combinations of the non-deleterious alleles and survive. However, we detected a moderate increase in medium and long ROHs in the East Pacific population over time, indicating more frequent mating among closely related individuals following the population decline. We also observed unusually wide variation in ROH patterns among individuals in the Northeast Caribbean, where some show few but extremely long ROHs, while others display ROH profiles similar to other small-stable populations. Such high heterogeneity in breeding between close relatives has been suggested as an early warning sign in small and/or declining populations where the chances of finding a non-relative mate decrease ^72^. This could also be influenced by other factors related to sea turtle life history traits, such as natal homing behaviors and breeding system dynamics. For example, there is documented high male competition for mates and multiple paternity and breeding sex ratios appear to be balanced in most leatherback populations even despite apparent female-biased hatchling sex ratios ^73^. This suggests that currently male scarcity is not an issue, but is a potential concern under climate warming because sea turtles have temperature dependent sex determination ^74^ where males may become more limited in the future. Further genetic monitoring with greater sample sizes across life stages is needed to robustly assess these dynamics, and to understand possible role of viability selection in ROH signatures and inbreeding depression as has been observed in other species ^8,75^. Collectively, interpreting leatherback genomic indicators in the context of their historical and contemporary demography offers hope for population resilience, but clearly warrants continued monitoring to detect and mitigate possible future genomic erosion and inbreeding depression.

The majority of leatherback populations are currently threatened with extinction, including in the Northwest Atlantic where populations were apparently generally stable, even increasing slightly, through 2010, but have declined significantly in recent and longer-term at site- and regional scales ^44^. The lack of recovery despite decades of conservation efforts has been perplexing, with possible contributions of genetic factors hypothesized but unknown. To this end, our study provides several key insights. Persistently low genomic diversity relative to other sea turtles resulting from long-term small effective population sizes consistent with previous estimates ^34,76^ demonstrates the species’ ability to persist over thousands of years with very limited genetic variation. This could provide resilience to contemporary population fluctuations, and may contribute to the ability of formerly small populations in the Northwest Atlantic to rapidly increase in abundance recently ^44^, and the persistence of some Pacific leatherback populations that were predicted to go extinct over twenty years ago ^26^. Nonetheless, their history doesn’t necessarily predict their future ^2^. On the one hand, leatherback evolutionary history facilitates effective purging of highly deleterious mutations to confer resilience under contemporary population declines. On the other hand, increased inbreeding and associated genetic load can erode adaptive potential and hinder recovery, particularly in the smallest populations ^3,77,78^ and under synergistic threats and rapid environmental change. A major remaining gap in our understanding of risk and resilience in leatherback turtles is linking genomic health metrics to fitness data (i.e, survival and reproductive output), to more fully understand how these indicators connect to inbreeding depression and population dynamics. Individual variation in reproductive fitness linked to maternal identity is a key driver of leatherback hatching success, which is lower and much more variable compared to other sea turtles ^79,80^. In addition to environmental influences and resource allocation, this may likely be connected to their genomic health and potentially a key explanatory factor for why they have lagged behind other species in their recoveries, but has not been investigated. Our findings lay the groundwork for urgently needed additional studies across leatherback populations and the sea turtle clade (similar to ^8,16,64^) to more fully understand if hatching success or other fitness measures are linked to inbreeding depression, low genetic variation, and/or other genomic drivers.

### Conclusion

Our study provides the first in-depth analysis of genomic health of leatherback sea turtles at a critical time as effective conservation strategies are needed to reverse declines and support long-term sustainable populations. Our findings suggest that despite low genomic diversity and small contemporary population sizes, the demographic history and life history traits of leatherback turtles likely provide resilience to support the recovery of healthy leatherback populations as long as current anthropogenic threats are effectively mitigated. Our results can provide the foundation for future research linking genomic health, fitness and population dynamics, monitoring genetic erosion over time, and examining genomic health in missing or sparsely sampled leatherback populations in this study. Beyond sea turtles, this work provides a clear demonstration that genomic and abundance-based metrics are complementary and can be used in tandem to assess extinction risk and guide conservation efforts, and also highlights the high importance of biobanking, long-term population monitoring programs and global collaboratives to robustly address these questions in wild populations of conservation concern.

## Methods

### Demographic metrics reconstruction and classification

We grouped leatherback nesting rookeries into populations following best available knowledge of demographic isolation and connectivity from prior studies ^42^ ^81–83^, confirming with population genetic analyses of our samples (see details below). We note that there are multiple types of population units defined for sea turtles used in management frameworks for different conservation goals (e.g., regional management units, distinct population segments, management units, genetic stocks; see ^84^ and ^85^ for details). It is most appropriate for the purposes of the research questions in this study to define populations as those with evidence of genetic distinction and demographic isolation from one another that are expected to be on independent evolutionary trajectories. Generally, these population groupings align with established “management units” (MUs), with some modifications based on fine-scale genomic analyses here or recent companionate information. For each sample (from individual turtles), we then followed the criteria for scoring population abundance and short-term trend following Wallace et al. 2011 and 2025 ^42,86^ over a ∼10-year period. We supplemented and validated these criteria with abundance-based metrics (i.e., nesting females and/or nests) and trends extrapolated from global and regional leatherback assessment reports ^22,42,44,49^. Samples were scored based on collection year, and for any sample that was taken prior to 2001 (i.e., outside of the time horizon for ^86^) we followed their original procedure to assign the abundance and short-term trend scores. Finally, to simplify our groupings for statistical analyses, we adapted abundance and trend categories from a discrete numerical scale to categorical. For abundance, we applied a simple threshold to the Wallace et al. 2025 ^42^ scoring definitions, where scores of 1 to 2 were designated as ‘large’ and 2.5 or 3 as ‘small.’ For trends, we combined the Wallace et al. 2025 ^42^ scoring definitions (1 = increasing, 2 = stable, 3 = decreasing) with data on prior population size to categorize as recovering, stable, or declining.

### Sample collection and DNA extraction

All samples were taken from reproductively mature females except where noted otherwise in Table S1 or S2. We obtained tissue (skin, blood or muscle) or DNA samples from the Marine Mammal and Sea Turtle Research (MMASTR) collection at NOAA Southwest Fisheries Science Center, collected from 1992-2023 (Table S1) and stored at −20°C or −80°C. All genomic DNA (gDNA) used in this study had been either previously isolated from subsamples of these tissues, or were extracted during the course of this study using one of the standard extraction techniques as described in ^43^. We assessed DNA integrity for samples used as input into both WGR and Rapture libraries by using a Fragment Analyser (Agilent, Santa Clara, California) or 4200 TapeStation System (Agilent, Santa Clara, CA), and/or quantified DNA concentration using a Qubit 4.0 Fluorometer (Invitrogen, Waltham, Massachusetts) following the recommended protocols provided by the manufacturers. After extraction, gDNA was stored at −80°C until use in downstream analyses.

### Library preparation and sequencing

For WGR data, we prepared libraries for N = 167 individuals using an Illumina Nextera PCR-based protocol adjusted for one tenth reaction sizes, as described in ^34^, adapted from ^87^. We used a Fragment Analyzer (Agilent, Santa Clara, CA, USA) to assess library quality, followed by pooling based on normalized concentrations and sequenced at Novogene (Sacramento, CA, USA; NovaSeq 6000) using 150-bp PE sequencing chemistry. A subset of the data for these samples were previously published in ^88^. We then supplemented this initial dataset with additional samples sequenced in companion projects: (1) N = 42, libraries were prepared by Novogene, Inc. (Sacramento, CA) using a standard PCR-based library preparation protocol with an average insert size of approximately 350 bp, followed by sequencing 150 bp paired-end reads on an Illumina NovaSeq X Plus platform, (2) a single library constructed and sequenced on an Illumina HiSeq 3000 platform (150 bp paired-end reads) at the University of Florida’s Interdisciplinary Center for Biotechnology Research Core NextGen Sequencing Core, and (3) N = 28 libraries constructed and sequenced by Element Biosciences (San Diego, CA, USA), as described by ^89^. We performed downstream checks to detect and account for any batch effects on data due to these differences in laboratory processing; combined, we generated WGR data for a total of N = 238 individuals (Table S1). The overview of the Rapture samples can be found in Table S2. Full details for Rapture libraries and sequencing are described in ^43^.

### Data pre-processing and variant calling

All code used for analyses is available in the corresponding github repositories listed in the Data Availability section. For WGR data, we assessed raw data quality using Fastqc (v0.11.9) ^90^ before and after trimming sequences for quality and adaptor contamination using bbduk (v38.90) ^91^. We aligned reads to the *Dermochelys coriacea* reference genome (NCBI accession GCA_009764565.4) ^34^ using BWA-mem with default parameters (v0.7.17) ^92^. We estimated mapping percentages, sorted and indexed files using SAMtools (v1.14) ^93^. We assessed mean depth using Mosdepth ^94^ and marked duplicates using markduplicates in Picard with default settings (v2.26.2) ^94^. We reassessed depth and read loss post duplicate removal, then re-aligned bam files around indels using GATK (v4.2.3.0) ^95^. We filtered out samples with <8X mean coverage and/or <95% mapping rate, and used GATK to call single nucleotide polymorphisms (SNPs) adapted from GATK best practices for workflow and filtering ^96^. Briefly, we called variants with HaplotypeCaller for each sample, imported individual gvcf’s into GenomicsDB prior to performing JointGenotyping and selecting variants, followed by filtering variants (--min-meanDP 5 --max-meanDP 200 --mac 1) using VCFtools (v0.1.14) ^97^. Following all filtering steps, we retained N = 222 samples and 13,590,825 variants; among those, 9,678,182 were SNPs. To remove any closely related or duplicate samples, pairwise relatedness between individuals was estimated using the full set of WGR variants. To achieve this, we converted the VCF file PLINK format, and generated relatedness metrics leveraging the inbuilt KING relatedness estimator (--make-king-table) ^98^. Pairwise matrix of kinship coefficients for all WGR leatherback samples can be found in Table S8.

Rapture sequencing data were demultiplexed, filtered to retain only samples with >10,000 sequences, and pre-processed as described in ^43^. We estimated coverage with bedtools, followed by using GATK to call SNPs following best practices as described above. We used VCFtools to filter variants (minimum mean depth = 5, max mean depth = 200, minor allele count = 1), then BCFtools ^93^ to concatenate filtered VCFs per chromosome into a single VCF. We imported the VCF file into R and further filtered to retain SNPs (min mean depth = 20; max mean depth = 200; per genotype min depth = 8). We then ran an iterative data missingness filter per individual and locus to optimize the number of IDs and loci retained, followed by pruning for linkage disequilibrium (-m 0.2 -w 1000) using BCFtools +prune plugin function ^93^. Finally, we used PLINK ^99^ to calculate relatedness metrics to identify any duplicate or highly related individuals. We removed 1 sample from pairs with high kinship values based on Z1 and Z2 scores following cutoffs as recommended in ^98^ and retained samples collected only from nesting turtles for a final dataset of N = 636, SNP loci = 4683.

### Population structure analysis

We used PLINK (v1.90p)^99^ to convert the final whole genome VCF of 222 leatherback individuals into BED format. We next filtered out sites that had an association r2 value greater than 0.2 in sliding windows of 50 SNPs (--indep-pairwise 50 10 0.2), resulting in 6,345,315 putatively unlinked SNPs. We note that leatherback turtles can have complex patterns of LD ^89^, so we also tested more conservative LD filtering thresholds as well as running analyses without LD pruning, and observed no substantial differences in the results. We next ran PLINK PCA and visualized the results in python3. To better resolve population structure among Northwest Atlantic rookeries, we additionally ran PCA analysis separately on samples belonging to Northern, Northwest, Northeast, and Southeast Caribbean locations (see location details in Table S1). To analyze population structure, we next performed ADMIXTURE (v1.3.0) ^100^ analysis varying the number of ancestral populations (K) from 2 to 8. We ran admixture analysis three times: 1) on the VCF with all 222 leatherback samples; 2) on the VCF with the samples belonging to the Eastern Pacific population; and 3) on the VCF with samples belonging to the Western Pacific and Indo-Western Pacific populations. The additional analyses on Eastern and Western Pacific populations separately allowed us to look into finer population structure in these groups. We selected the K value with the lowest cross-validation error. We visualized ADMIXTURE results in python3.

For the Rapture data, we converted the VCF of 636 leatherback individuals to a genlight object (package adegenet) and genind object (package dartR) ^101,102^ to analyze Fst and bootstrapped Fst p-values among nesting sites and populations, calculate principal component scores, and run a discriminant analysis of principal components (DAPC) to estimate the optimal number of clusters represented in the data. We filtered out sites that have an association r2 value greater than 0.1 in sliding windows of 50 SNPs (--indep-pairwise 50 10 0.1). We conducted DAPC analyses ^103^ for the entire dataset (n = 636), as well as for only the Pacific basin (n = 313), Atlantic basin (n = 306), Western Atlantic (n = 215), and Western Pacific (n = 236). We complemented these analyses with ADMIXTURE ^100^ to estimate the number of ancestral populations within the sites sampled in the Western Pacific (K = 2 - 7). Visualization was done with the pophelper R package ^104^.

### Demographic modeling

The remaining analyses focused on whole genome data. Datasets used for demographic reconstruction were filtered solely for quality as described in the previous sections and not for LD or MAF. To reconstruct historical demography, we first applied the Pairwise Sequentially Markovian Coalescent (PSMC) model (v0.6.5) ^105^ to a high-coverage representative individual from each population. Diploid autosomal consensus sequences were generated using BCFtools consensus against the leatherback turtle reference genome ^34^. We utilized a generation time of 30 years and a mutation rate of 1.2e-08 following ^34^, with the recommended command structure (p = 4+25*2+4+6). We performed 100 bootstraps (split by chromosome for efficiency) to assess variance. We then estimated recent effective population size (Nₑ) separately for each population using GONE2 (Genetic Optimization for Ne Estimation) ^106^, which infers Nₑ from linkage disequilibrium (LD) patterns in whole-genome SNP data. We included 171 samples (Table S1). For each population, we extracted biallelic SNPs from a joint VCF file using VCFtools (v0.1.15)^97^ and removed sites containing indels. Filtered data were converted to PLINK format (.ped and .map) using PLINK (v1.9) ^99^. We calculated sex-averaged genetic distances (cM) for each SNP using chromosome-specific recombination rates ^89^, scaling positions by chromosome length following ^107^. To test the robustness of GONE2 estimates, we conducted a separate leave one out analysis, in which one random individual was excluded from each of up to four runs or until every individual had been removed once for smaller sample sizes.

### Population trend modeling

To estimate annual population trends, we implemented a Bayesian State-Space Model using the jagsUI package in R ^48^. Population abundance data (all based on nests, except tracks for Southeast Atlantic/West Africa) ^22^ were log-transformed to stabilize variance and modeled as multivariate time series. The model partitioned data into a hidden state process, representing the “true” population trajectory, and an observation process, accounting for sampling error across multiple time series within a single population. The state process followed a random walk with a constant drift term (U), representing the mean growth rate and process error variance (Q). Observation error was estimated independently for each time series. We ran three MCMC chains for 20,000 iterations each, with a 5,000-iteration burn-in and a thinning rate of 5. Posterior distributions of total abundance were back-transformed to the arithmetic scale, and a LOESS regression (span = 0.9) was applied to the resulting medians and 95% symmetric credible intervals to produce a smoothed population trend for visualization.martin

### Runs of homozygosity estimation

To estimate runs of homozygosity (ROH), we used PLINK (v1.90p) ^99^. This method employs a rule-based approach that can be sensitive to parameter settings, depth of coverage and demographic history ^69,108^. Therefore, we first conducted preliminary scans of the relationship of depth of coverage on ROH detection across and within samples (via random down-sampling), confirming that an 8X filtering threshold was effective in removing technical artifacts and aligning with prior literature ^69^. Secondly, to optimize parameters to be most appropriate for leatherback turtles that have low genomic diversity and slow mutation rates ^38^, we allowed a lower minimum of 20 SNPs as in ^34^, and tested SNP sliding window size (--homozyg-window-snp) from 10 to 200 with step 10. To estimate how this parameter impacted recovering ROHs of different lengths, we binned ROHs into four bins of sizes between 50 and 100 kb, between 100 and 500 kb, between 500 and 1000 kb, and above 1000 kb. These analyses confirmed larger SNP sliding window sizes decreased the number of detected ROHs. Given the conservation context of this study, we prioritized avoiding false negatives over false positives (detecting more ROHs rather than missing real ROHs). Accordingly, we applied a more conservative approach and ran the final analyses with the following parameters: --homozyg-snp 20 --homozyg-kb 50 --homozyg-window-snp 20 --homozyg-window-het 1 --homozyg-window-missing 5 --homozyg-window-threshold 0.01. Inbreeding coefficient (F_ROH_) was estimated as the total length of ROH / genome size, where the genome size was the size of the reference leatherback genome assembly (2,164,762,090 bp). ROH size classifications can vary widely across studies with little justification. While most use thresholds for long ROH of >1Mb to >10Mb in mammalian systems assuming recombination rates of ∼1cM/Mb, it has been suggested that for other taxonomic groups with higher recombination rates, lower thresholds may be more appropriate ^13^. Recombination rates for leatherback turtles substantially vary across their genome from 0.11 - 2.8 cM/Mb ^89^, we therefore applied a similarly conservative approach as above for defining ROH categories as: small =<100 kb, medium = 100-500 kb, long =>500kb. Finally, we re-confirmed that our ability to recover ROHs was not affected by coverage, as indicated by the lack of correlation between any of the ROH metrics (total number, the aggregate length, or the average length of ROHs), including ROH separated into length classes, and the resequencing depth (Fig. S5).

### Genetic load analysis

To assess the genetic load in the study populations, we employed a combination of annotation-based and conservation-based approaches using two primary tools: snpEff and ANNOVAR. Below, we outline the workflow for variant annotation, classification, and analysis using both methods.

Variant annotation was performed using snpEff (v2017-11-24) ^109^ to classify variants by their predicted functional impact (HIGH, MODERATE, LOW, MODIFIER). The input included a combined VCF file, a reference genome, and a GTF annotation file. Custom genome and annotation databases were configured in the snpEff.config file. The annotated VCF was split by impact category using BCFtools (v1.19) ^93^, and further subdivided by sample and genotype (heterozygous/homozygous). Counts of SNPs and indels per impact category were aggregated using custom scripts (count_var_by_impact.sh).

For cross-validation, variants were re-annotated using ANNOVAR ^110^. A custom database (turtledb) was generated from the genome annotation. Variants were classified into functional categories (e.g., frameshift, stopgain, synonymous) and mapped to snpEff’s impact levels using a predefined correspondence: frameshift variants, stopgain/loss in ANNOVAR were annotated as high-impact variants; nonframeshift, nonsynonymous SNVs in ANNOVAR - as moderate-impact variants; synonymous SNVs in ANNOVAR - as low-impact variants; and unknown/other variants in ANNOVAR - as modifier variants. Filtering and splitting followed the same workflow as snpEff.

We estimated relative enrichment of derived (non-reference) alleles (Rxy) between larger-declining and small-stable populations using a block jackknife approach. An annotated SNP VCF was split by demographic trend (small-stable and large declining) and by predicted functional impact class (HIGH, MODERATE, LOW, and MODIFIER). For each impact class and population group, we extracted genotypes, converted them to derived allele counts per site, and divided the resulting site-wise counts into 100 contiguous blocks. To obtain jackknife replicates, we iteratively recalculated the total derived allele count while leaving out one block at a time, for both larger-declining and small-stable populations. For each impact class, we then computed Rxy as the ratio of derived allele counts in the two populations, scaled by their respective sample sizes. Finally, Rxy values for HIGH, MODERATE, and LOW impact categories were normalized by the corresponding Rxy estimated for MODIFIER sites.

### Temporal analysis

To evaluate temporal changes in heterozygosity, inbreeding, and genetic load across the Northeast Caribbean and Eastern Pacific populations, we modeled each parameter using generalized linear models with generation as the main predictor of a trend. For the aggregate length of Runs of Homozygosity (ROHs) and genetic load metrics we used a gamma distribution and for the number of ROHs we used a negative binomial distribution. We analyzed the inbreeding coefficient (F_ROH_) and heterozygosity using a logit-transformed linear model to accommodate their bounded 0-1 distribution. P-values for ROH and genetic load metrics were corrected for multiple testing using the Benjamini-Hochberg false discovery rate (FDR) procedure. In the figures, we report FDR-corrected p-values together with the direction of the trend (slope sign) when the trend was significant (FDR-corrected p < 0.05).

### Structural variation analysis

To assess large-scale structural variations between groups across each chromosome, we used a window-based local PCA approach using the WinPCA python package (v1.2) ^111^. We initially subsetted the final VCF file of all 170 individuals for each chromosome with VCFtools (v0.1.16) ^97^, and subsequently extracted only bi-allelic SNPs using BCFtools (v1.21) (--types snps -m 2 -M 2)^112^. For each chromosome (1-28), we calculated PCAs using 1 Mb sliding windows, with a 100 Kb interval across all groups. The ‘chromplot’ function of WinPCA was run, using the pre-determined regional management groupings, and plotting every 10^th^ PCA interval to visualize differences between groups. The resulting HTML files of PC1 were screened by eye to identify regions that showed patterns of divergence between one or more of the regional management groups, and that showed higher percentages of variance explained by PC1 (e.g., inversions, characterized by a splitting of all individuals into three distinct groupings on PC1). From all of these regions of interest, two genomic regions were selected for deeper investigation. We extracted the functional annotation from these regions using the leatherback turtle RefSeq annotation (GCF_009764565.3; Annotation Release 101). Gene symbols, gene names, and (in instances where functional annotation was insufficient to assign these) protein annotations were used to determine the functional significance of these divergent areas. To assess the genomic compatibility of the putatively inverted region in chromosome 14, the WinPCA analysis was applied to a subset of individuals from the dataset with known kinship, because only compatible genotypes would be predicted to be observed in parents or offspring that have completed embryonic development. Data were available for a subset of mother-offspring relationships from Mexico (Eastern Pacific population), and grandmother-mother-offspring from St. Croix (Northeast Caribbean population) relationships. Incompatible genotypes are not expected to appear in the PCA, with the presence of heterozygotes indicating genomic compatibility at least through embryonic development in these putative inversions.

## Acknowledgements

The authors would like to thank the many people and organizations working on leatherback nesting beach research, monitoring and conservation programs that made this research possible, especially John Pita, Simon Vuto and Peter Waldie in the Solomon Islands, Neca Marcovaldi and Fundação Projeto Tamar, Rachel Thomas and the University of Florida Sea Turtle Hospital, and the Gabon Agence Nationale des Parcs Nationaux and their national partners and field teams. The authors would also like to thank Reid Brennan for helpful comments on an earlier version of this manuscript. We thank Gabriela Serra-Valente, Alaina Harmon, Victoria Pease, Suzanne Roden, Alexandria Mena and Dan Prosperi for sample curation and processing at NOAA Southwest Fisheries Science Center. Bioinformatic analyses were conducted using the UMass UNITY supercomputer supported by the Massachusetts Green High Performance Computing Center (MGHPCC). This research was supported by funding and support from the University of Massachusetts Amherst, National Science Foundation IOS (grant #1904439 to L.M.K.), NOAA-Fisheries, the United States Fish and Wildlife Service Marine Turtle Conservation Fund, Element Biosciences, The Sea Turtle Conservancy (Florida Sea Turtle Grants Program project number 17-033R to D.J.D.), and the National Save The Sea Turtle Foundation (the Fibropapillomatosis Training and Research Initiative to D.J.D.).

## Data and Code Availability

Population resequencing data of all leatherback sea turtle individuals generated in this project are available at NCBI BioProjectID PRJNA1446838 (whole genome resequencing data) and PRJNA1454575 (rapture sequencing data), which will be made public upon publication. The code for all analyses is available at https://github.com/osipovarev/Leatherback_popgen, https://github.com/kfphillips/leatherback_rapture/, and https://github.com/sebasalco/LeatherbackTurtles_Demographic_History.

