## Supplementary Figures for "Genomic indicators of risk and resilience in global leatherback turtle populations"

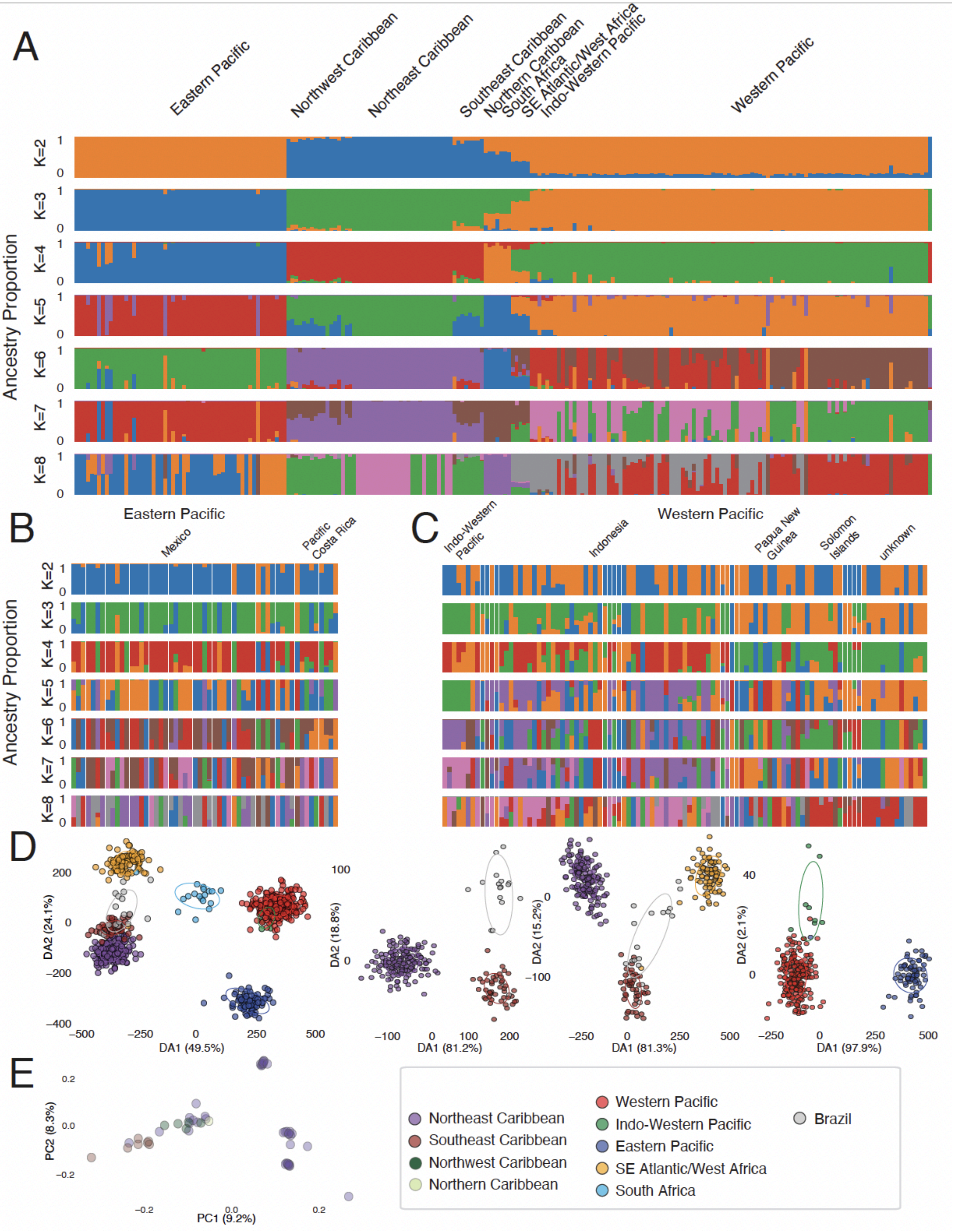

**Fig. S1: Population structure analysis of leatherback samples.**
(A) Admixture analysis of all whole-genome samples (n = 222), with the number of ancestral populations (K) ranging from 2 to 8. Black lines above each plot indicate individual locations where samples were collected. Colored lines above the entire panel indicate populations.
(B) Admixture analysis of Eastern Pacific population individuals (K = 2 - 8).
(C) Admixture analysis of Western Pacific and Indo-Western Pacific populations (K = 2 - 8).
(D) DAPC analysis of Rapture data. The left-most plot shows all samples (n = 636); subsequent plots show analyses of selected groups to improve resolution among populations.
(E) PCA analysis of whole-genome resequencing data of Northwest Atlantic locations conducted to improve resolution among populations.

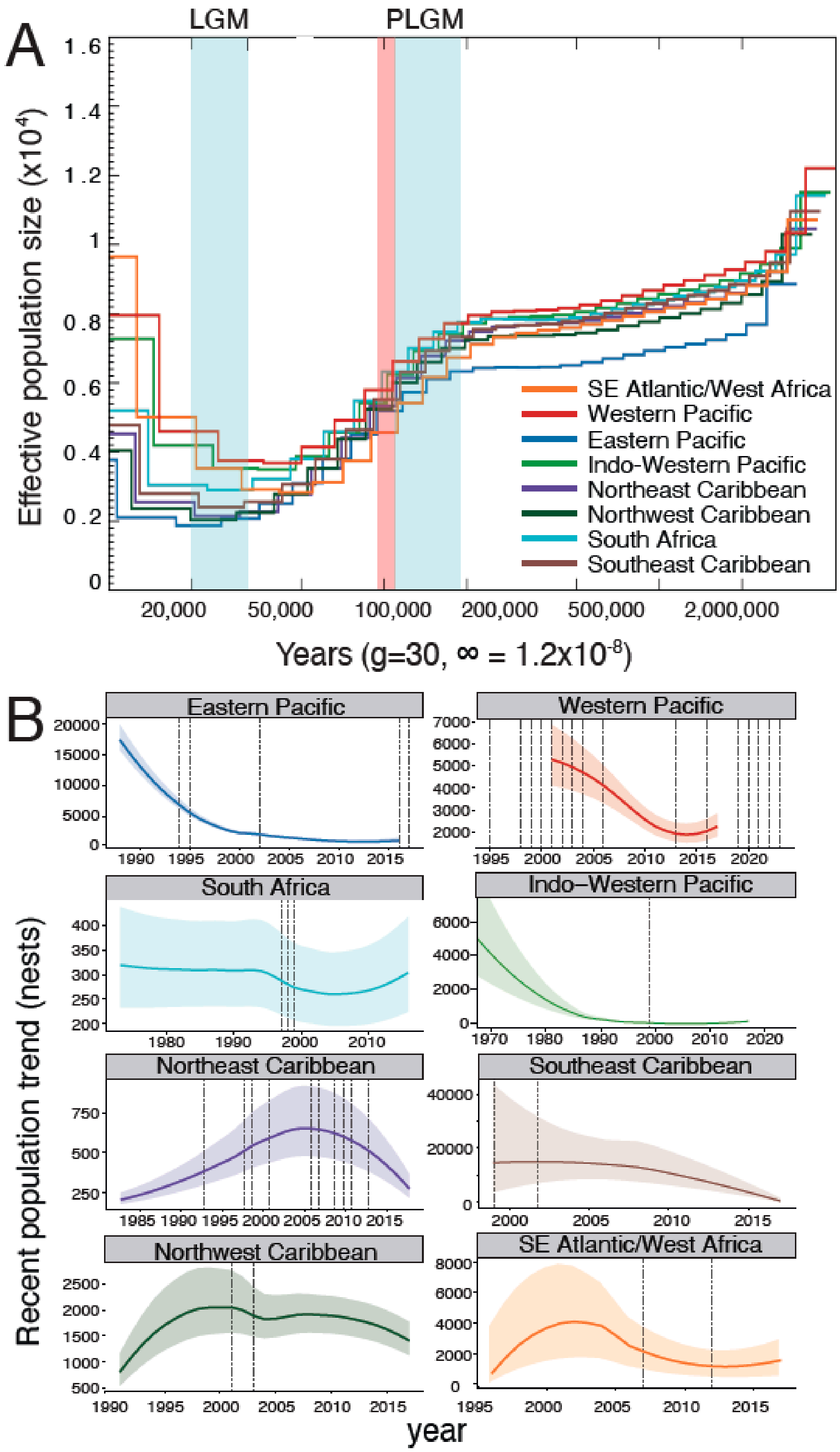


**Fig. S2: Demographic reconstructions of leatherback populations.**

(A) PSMC demographic reconstructions. All populations showed a gradual decline in Ne beginning around 2 million years ago until it reached an Ne of about 8,000 approximately 200,000 years ago. A more pronounced decline began for all populations 180,000 - 120,000 years ago, coinciding with the Penultimate Glaciation, and continued until the onset of Last

Glacial Maximum (LGM, around 26,000 years ago). Following the LGM all populations showed an increase in Ne, likely reflecting post-glacial population expansion and potential interoceanic gene flow facilitated by warming climates and changing oceanographic conditions, although the magnitude and timing differed between populations. Over these long evolutionary timescales, Ne estimates for all populations were relatively low: between 2,000 and 12,000.
(B) Recent population trends estimated from data collected between years 1967 and 2017 [48], with unconstrained axes to fully depict data for each population to facilitate evaluation of size and changes over time within each population. For example, this demonstrates the rapid declines of historically larger populations in the Indo-Western Pacific and Southeast Caribbean populations, the increases and decline in the smaller Northeast Caribbean population, and the steady stable trend in South Africa. Vertical dashed lines represent years when samples were collected for genomic analyses.

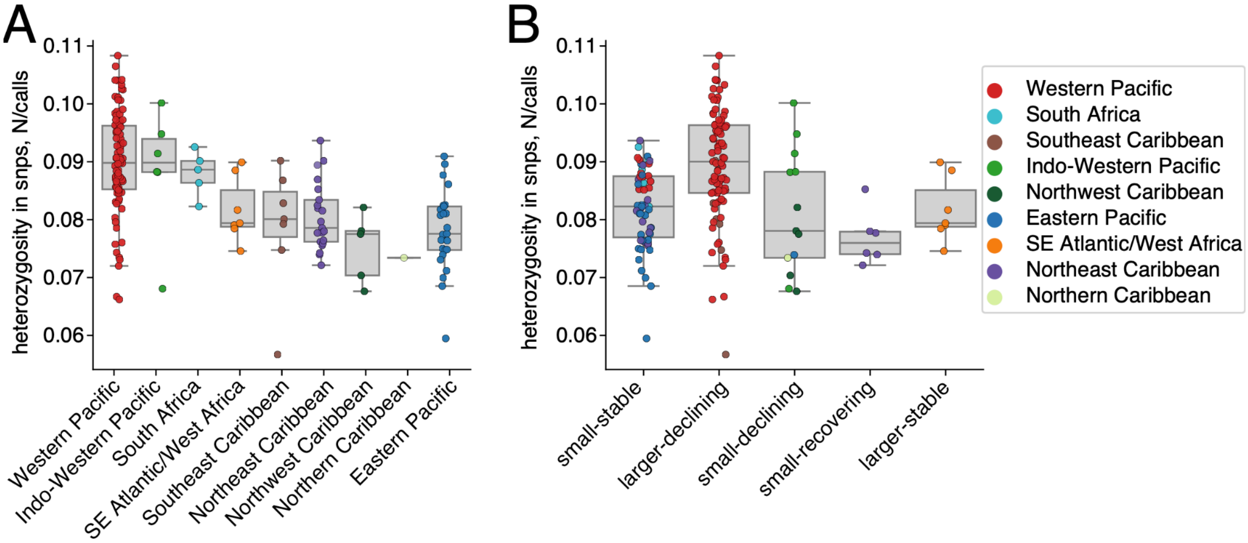


**Fig. S3: SNP density and heterozygosity across all leatherback populations.**
(A) heterozygosity presented for each population and (B) presented based on assigned demographic trends.

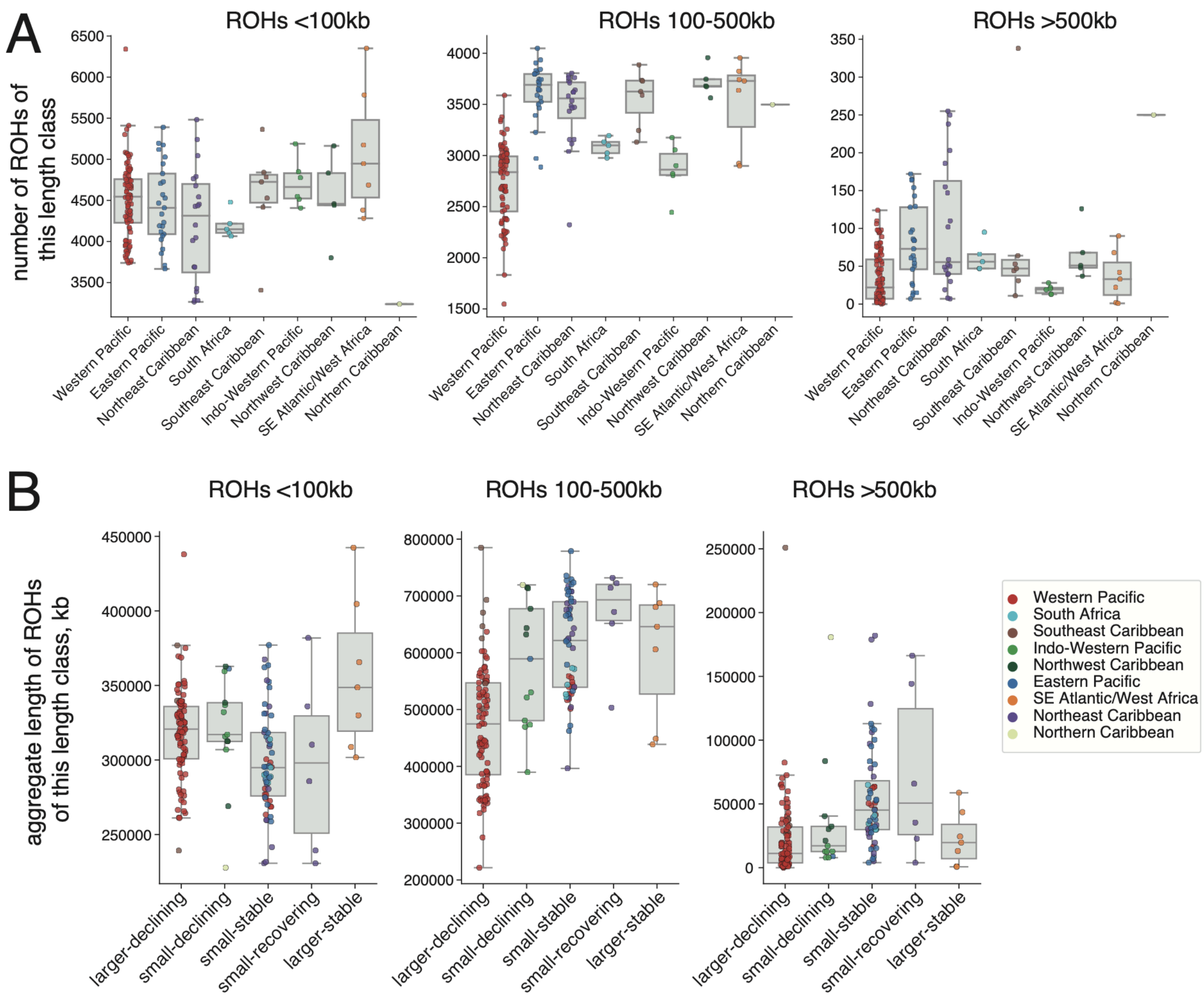


**Fig. S4: Number and length of Runs of Homozygosity (ROH) across all leatherback populations.** (A) Number of ROHs by length class. The number of medium-length ROHs (between 100 and 500 kb) shows the largest variability across leatherback populations.
(B) Aggregate length of ROHs by length class for each demographic trend.

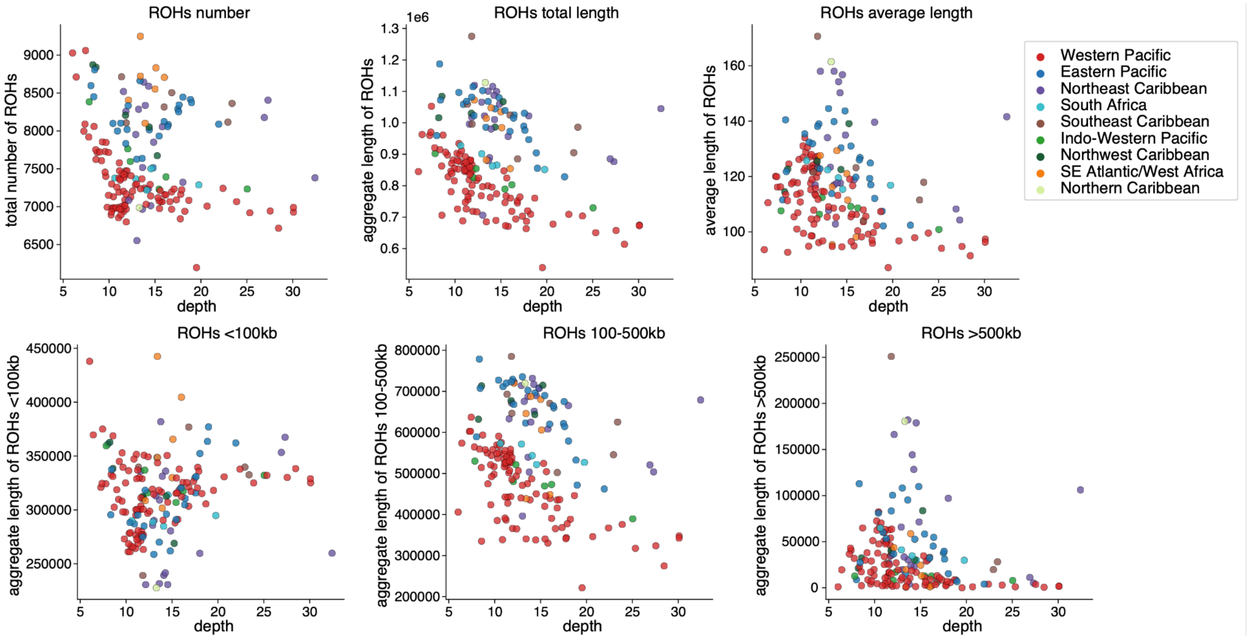


**Fig. S5: SNP depth does not affect the recovery of Runs of Homozygosity (ROHs).**
The total number, the aggregate length, or the average length of ROHs (in kb) show no correlation with SNP depth. ROHs separated into length classes show no correlation with SNP depth either.

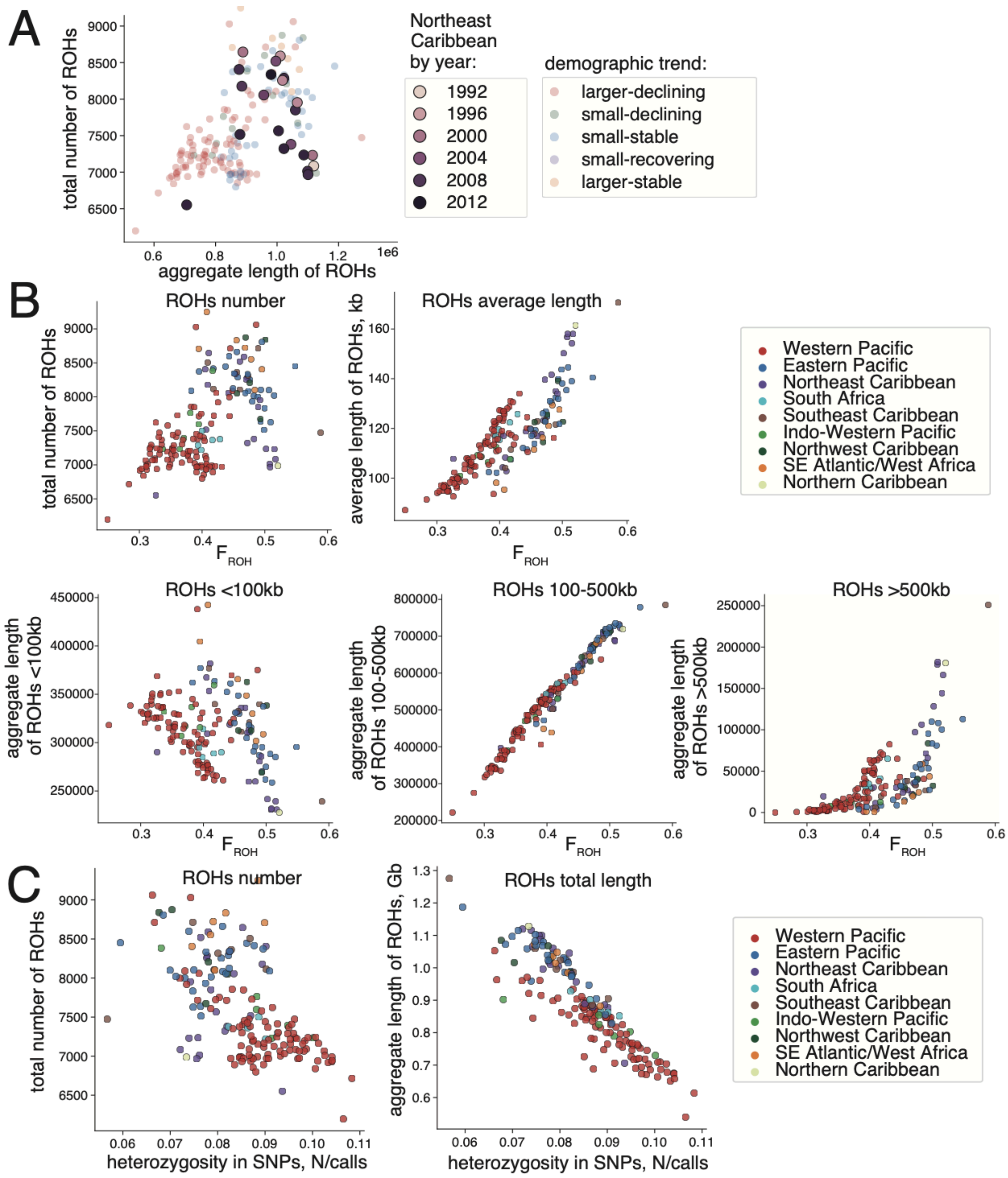


**Fig. S6: Runs of homozygosity (ROHs), inbreeding ($F_{ROH}$), and heterozygosity.**

(A) Distribution of ROHs in the North Caribbean population by year.

(B) The total number of ROHs, the total aggregate length of ROHs, the average ROH length, and the aggregate length of ROHs (in kb) separated by length class are presented as a function of inbreeding coefficient $F_{ROH}$. Patterns of inbreeding ($F_{ROH}$) in leatherback populations are driven primarily by medium-length ROHs (between 100 and 500kb).

(C) The total number of ROHs and the total aggregate length of ROHs are presented as a function of heterozygosity in SNPs.

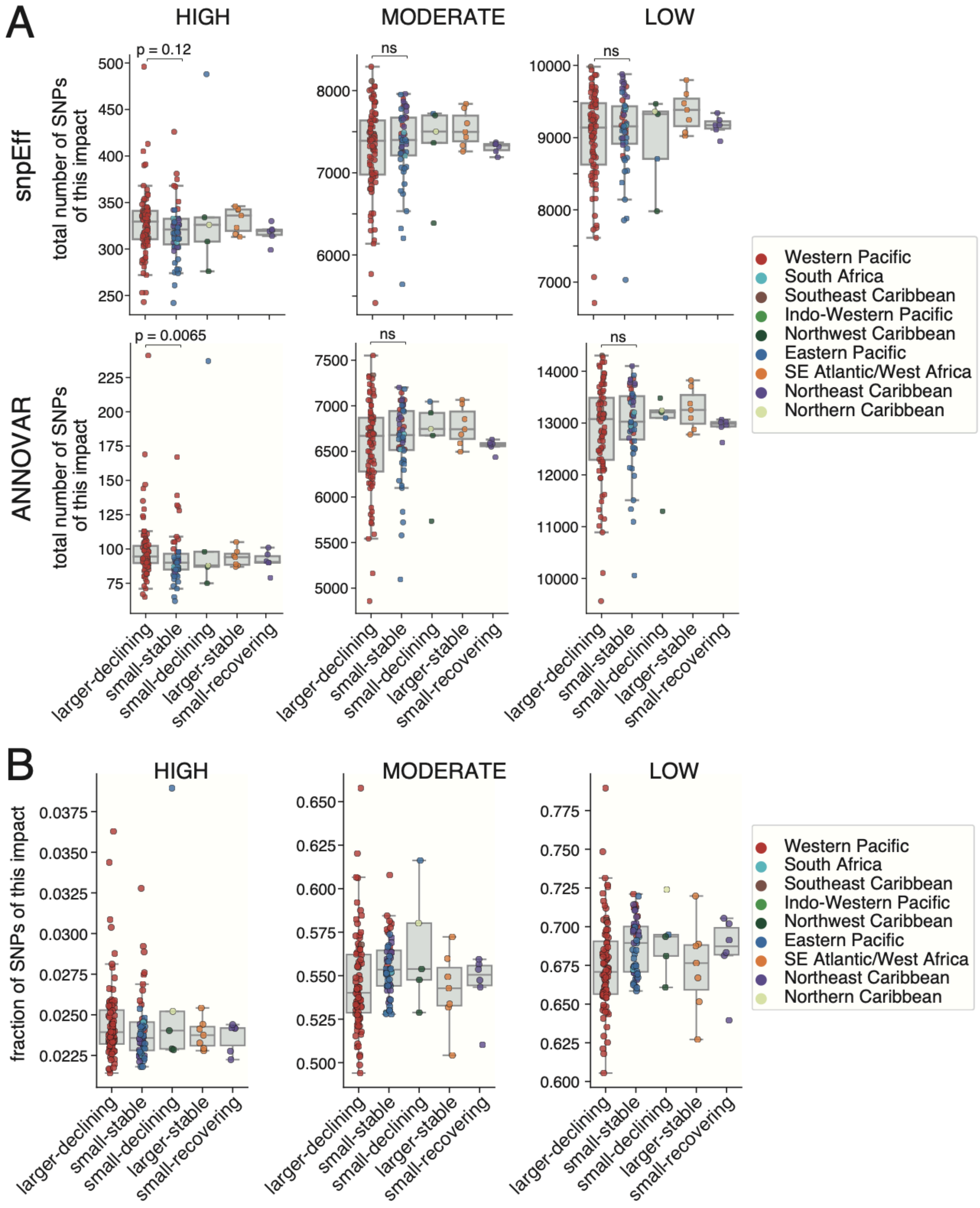


**Fig. S7: Total genetic load across leatherback populations based on demographic trend.**

(A) Total number of SNPs of each impact across all leatherback populations annotated with snpEff and ANNOVAR. For statistical comparisons between these two groups, a two-sided Mann-Whitney U test was used.
(B) Fraction of SNPs of each impact across all leatherback populations annotated with snpEff. 100% = high + moderate + low impact SNPs per individual; fraction of high-impact SNPs = high / high + moderate + low impact SNPs.

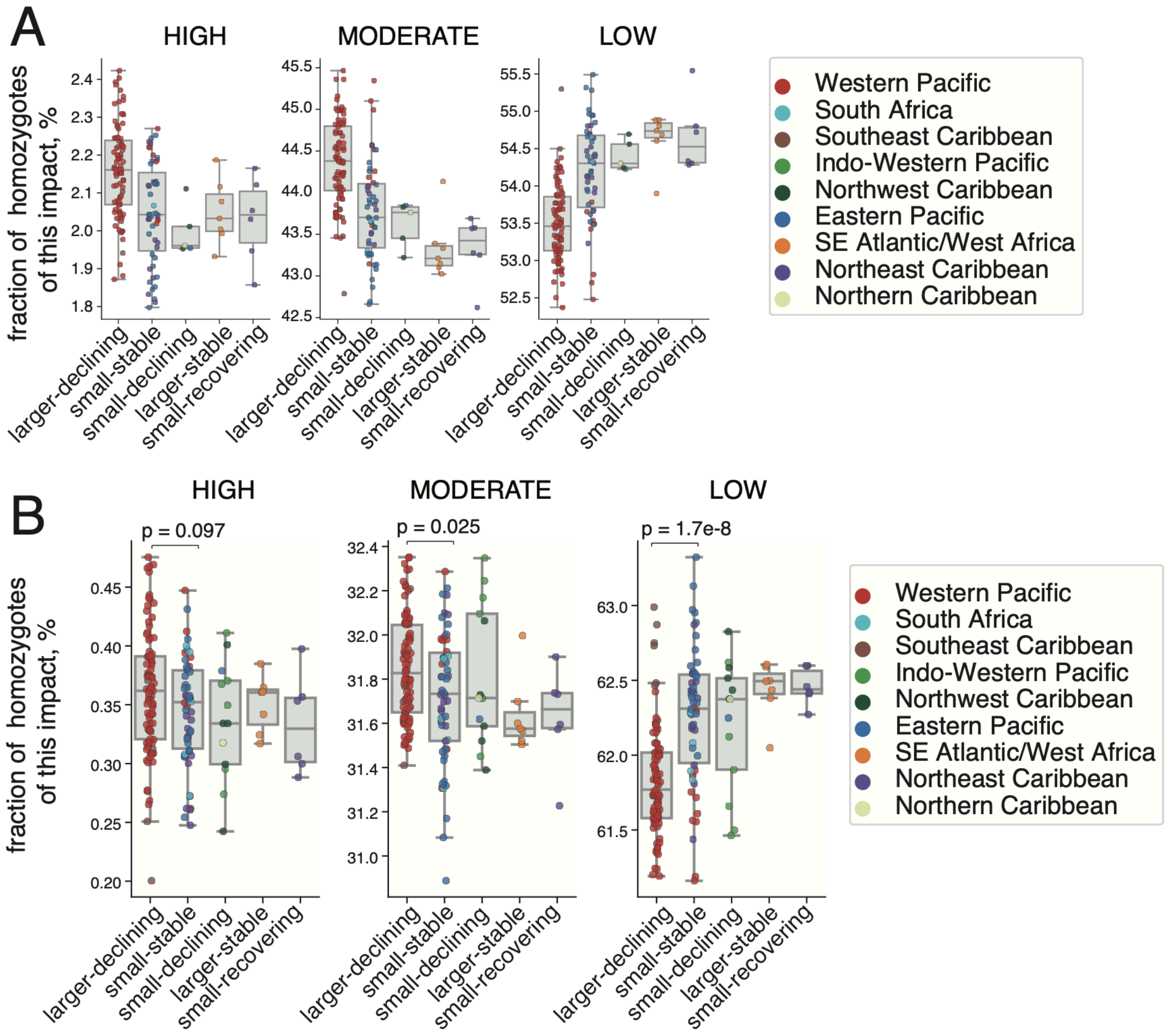


**Fig. S8: Realized genetic load based on homozygotes across all leatherback populations.**
(A) Fraction of homozygous SNPs of each impact annotated with snpEff.
(B) Fraction of homozygous SNPs of each impact annotated with ANNOVAR.
100% = high + moderate + low impact homozygous SNPs per individual; fraction of high-impact homozygous SNPs = high / high + moderate + low impact homozygous SNPs.

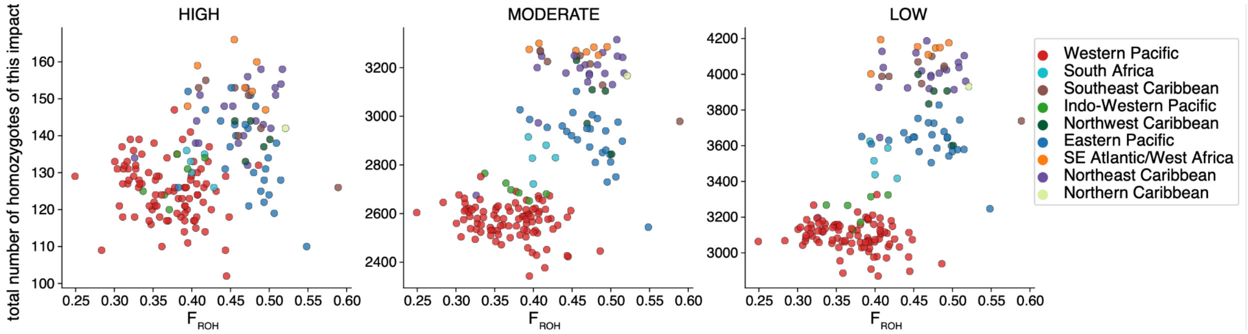


**Fig. S9: Number of homozygous SNPs per impact and inbreeding coefficient across all leatherback populations annotated with snpEff.**

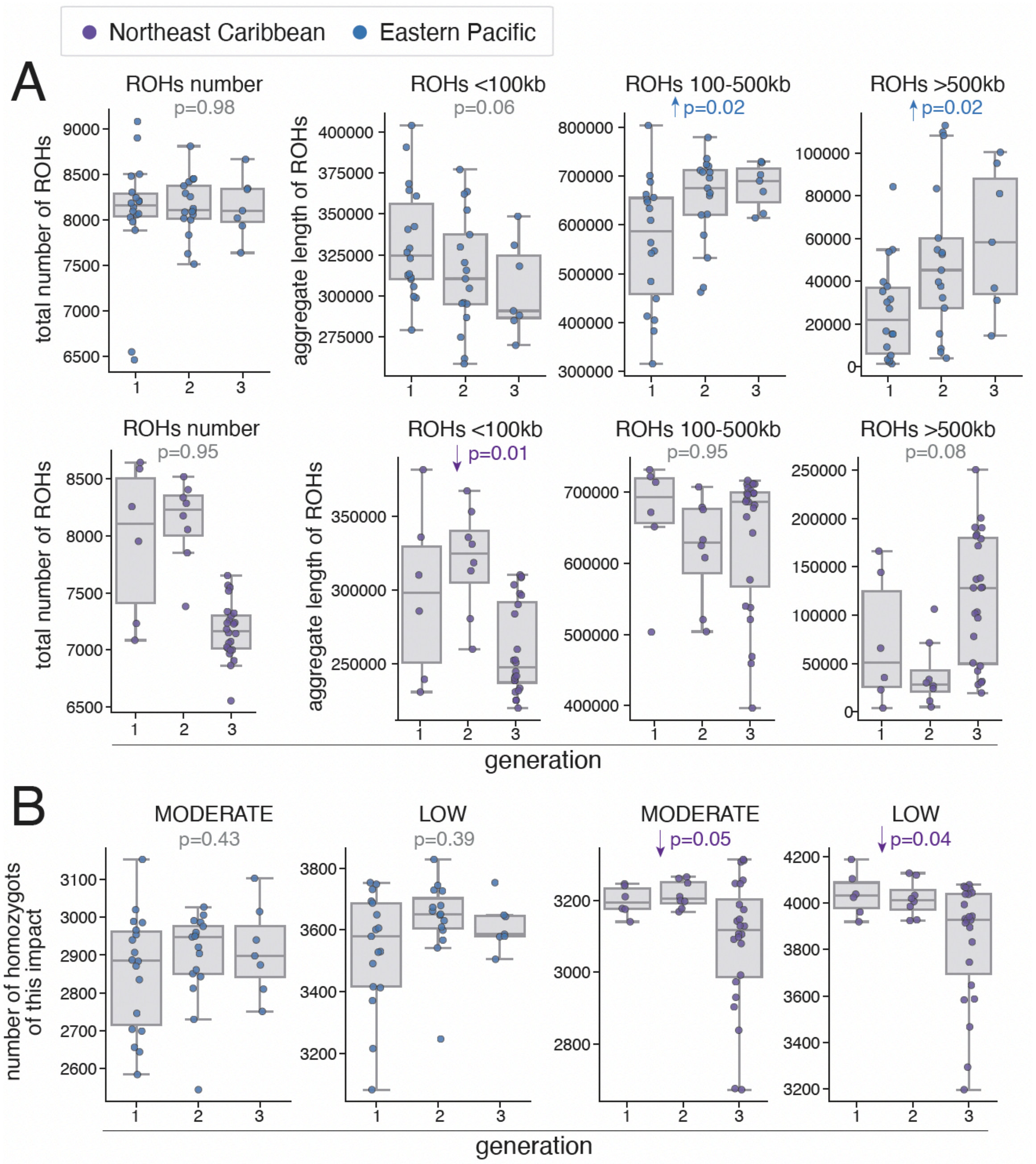


**Fig. S10: Temporal analysis of inbreeding and realized genetic load.**

(A) Inbreeding analysis based on Runs of Homozygosity (ROHs) over the past decades in the Northeast Caribbean and the Eastern Pacific populations.

(B) Genetic load measured based on the total number of homozygous SNPs by fitness impact over the past decades in the Northeast Caribbean and the Eastern Pacific populations. Variant fitness impacts are annotated with snpEff.

FDR-corrected p-values for the year effect are shown; p-value colors indicate significance, with grey denoting non-significant results ($P \geq 0.05$) and blue indicating a significant temporal trend ($P = 0.02$ for the Northeast Caribbean).
